# Task-induced modulations of neuronal activity along the auditory pathway

**DOI:** 10.1101/2020.07.11.198481

**Authors:** Gioia De Franceschi, Tania Rinaldi Barkat

## Abstract

Sensory processing varies depending on behavioral context. Here, we asked how task-engagement modulates neurons in the auditory system. We trained mice in a simple tone-detection task, and compared their neuronal activity during passive hearing and active listening. Electrophysiological extracellular recordings in the inferior colliculus, medial geniculate body, primary auditory cortex and anterior auditory field revealed widespread modulations across all regions and cortical layers, and in both putative regular and fast-spiking cortical neurons. Clustering analysis unveiled ten distinct modulation patterns that could either enhance or suppress neuronal activity. Task-engagement changed the tone-onset response in most neurons. Such modulations first emerged in subcortical areas, ruling out cortical feedback from primary auditory areas as the only mechanism underlying subcortical modulations. Half the neurons additionally displayed late modulations associated with licking, arousal or reward. Our results reveal the presence of functionally distinct subclasses of neurons, differentially sensitive to specific task-related variables but anatomically distributed along the auditory pathway.

## INTRODUCTION

Sensory perception is remarkably flexible and varies depending on the behavioral context. The response of sensory neurons can be modulated not only by external factors such as the statistics of the environment (Simoncelli and Olshausen, 2001), but also by the internal state of the animal, and the auditory system is no exception (Busse et al., 2017; Kuchibhotla and Bathellier, 2018; Poulet and Crochet, 2019). The activity of auditory neurons has indeed been shown to be influenced by several task-related variables such as attention or engagement (Atiani et al., 2009; Francis et al., 2018b; Fritz et al., 2003; Kato et al., 2015; Kuchibhotla et al., 2017; Otazu et al., 2009; Yao et al., 2019), arousal (Lin et al., 2019; McGinley et al., 2015), movement (Bigelow et al., 2019; McGinley et al., 2015; Nelson and Mooney, 2016; Nelson et al., 2013; Schneider et al., 2014b; Williamson et al., 2015; Zhou et al., 2014), or reward (Brosch et al., 2011; Gruters and Groh, 2012; Guo et al., 2019; Komura et al., 2001; Metzger et al., 2006). However, these task-induced modulations have been usually investigated separately, and it is not clear to what extent these signals are concurrently broadcasted throughout the auditory system or if they are selectively targeted to specific neural subpopulations. In this study, we investigate the functional and anatomical distribution of modulations associated with engagement, arousal, movement and reward along the auditory pathway.

The functional advantage of task-induced modulations is likely to be a representational improvement of behaviorally relevant stimuli. Previous studies on auditory engagement have mainly used spatial attention or discrimination tasks triggering selective attention, which is thought to enhance the discriminability of attended stimuli. The comparison of neural responses when the animals performed the tasks with passive hearing showed that the effects that selective attention has on auditory neurons are complex and heterogeneous. Specifically, it has been shown that attending to a specific target tone can induce enhancement or suppression of responses to the attended tone in both auditory cortex (ACx) and inferior colliculus (IC) (Francis et al., 2018b; Fritz et al., 2003; Fritz et al., 2005, 2007; Kuchibhotla et al., 2017; Otazu et al., 2009; Slee and David, 2015), that responses of cortical subclasses of inhibitory interneurons are differentially modulated (Kato et al., 2015; Kuchibhotla et al., 2017), and that cortical attentional shifts seem to be stronger in superficial layers (Francis et al., 2018a). The attentional modulation was also shown to be influenced by task difficulty (Atiani et al., 2009). However, the functional changes happening during non-selective attentional processes that require monitoring the whole auditory environment, such as detecting any sound, have been less investigated. It is not clear whether they share similar mechanisms with selective-attention processes. In addition to attentional load, task-engagement is likely to influence arousal levels. Arousal, generally assessed by measuring pupil size (Lin et al., 2019; McGinley et al., 2015), has been shown to affect both behavioral performance and sensory processing by improving the signal-to-noise ratio. Intermediate arousal levels correspond to optimal auditory tone-detection performance, and have been shown to be associated with changes in synaptic and circuit mechanisms that are stronger in the ACx than in the medial geniculate body of the thalamus (MGB) (McGinley et al., 2015). Elevated arousal has also been shown to improve the frequency discrimination of populations of neurons in superficial layers of the primary auditory cortex (A1) by broadening their frequency tuning while reducing noise correlations (Lin et al., 2019). If, and how, task-engagement and arousal differentially modulate auditory neurons is not clear.

Task-engagement is also often accompanied by movements not present in the passive state. Motor-related signals have been shown to be usually suppressive, robust and widespread across the auditory pathway (Schneider and Mooney, 2018). Auditory responses can be modulated by a variety of movements such as locomotion (Schneider et al., 2014a; Zhou et al., 2014), licking (Singla et al., 2017), vocalizations (Eliades and Wang, 2008), and other body movements (Rummell et al., 2016; Schneider et al., 2014a; Zhou et al., 2014). The effects that motor actions have on auditory responses have been shown to be stronger in cortex (Zhou et al., 2014), where they seem to be mediated by inhibitory neurons (Nelson et al., 2013; Schneider et al., 2014a). However, not all movement-related inputs are suppressive: small non-locomotor movements induce excitatory cholinergic activation of the ACx via basal forebrain inputs (Nelson and Mooney, 2016). It is not known if auditory neurons are only influenced by either engagement, arousal, movement, reward or by combinations of them.

Previous studies have clearly shown that the auditory pathway is not a simple feedforward network that exclusively processes sound signals, but rather a flexible system able to adjust responses depending on different behavioral contexts. Such modulations have usually been studied separately, focusing on either attentional, arousal, movement or reward effects on auditory responses in circumscribed brain regions. Here, we characterize the functional and anatomical distribution of distinct task-induced modulations along the auditory pathway. We performed extracellular recordings in the IC, medial geniculate body (MGB), and the two primary auditory cortices (A1 and anterior auditory field: AAF) of mice trained to perform a tone-detection task and compared neural responses in passive and active phases of the task. First, we show that most neurons along the pathway are modulated by task-engagement. We then unveil the presence of functionally distinct clusters of neurons, characterized by specific patterns of task-induced modulation reflecting either attention, arousal, movement, reward, either alone or in combinations. These clusters are present in all identified anatomical groups, but some are particularly represented in a specific region, cortical layer or neuronal subclass (putative regular or fast-spiking). Lastly, we show that modulations of onset auditory response to sound first emerge in the IC and MGB, ruling out feedback from ACx as the only mechanism underlying such subcortical modulations.

## RESULTS

### Mouse performance is reliably different between passive and active phases

To study how the internal state modulates responses in the auditory system, we trained 12 mice to perform a simple tone-detection task partitioned into passive and active blocks (Figure 1, see also Methods). Mice were head-fixed in front of a licking spout and first trained in the active phase, during which they had to lick the spout in response to a 1 s long pure tone within 2 s to receive a reward (Figure. 1a-b). Each active block consisted of 150, 65 dB SPL pure tones of randomly interleaved frequency (15 frequencies, logarithmically spaced from 4-45 kHz). To ensure engagement, the interstimulus interval (ISI) was randomly varied between 3 to 7 s and was reset if the animal licked outside the reward window. We tracked the animals’ training level by measuring their performance as the percentage of correct trials (% hit) and the average reaction time (Figure 1c, Figure S1). Once the mice performed well in the active phase of the task (A), we introduced a passive phase before (P1) and after (P2) the active block. The passive phases were identical to the active one, the only difference being the absence of reward in case of a hit (Figure 1b). The final task therefore consisted of a sequence of passive-active-passive (P1-A-P2) blocks. Animals were free to perform the task at any time, but trained mice were consistently engaged during the A-phase and disengaged during the P1 and P2 phases (Figure 1c-g, Figure S1). The passive and active phases of the tasks were also distinguishable from their hit response latencies, which were precisely time-locked to sound onset in the active phase but not in the passive one (Figure 1g-h). The clear differences in performance and hit latencies (Figure 1f-h) confirm that mice were in different engagement states during the passive and active phases.

**Figure 1.**
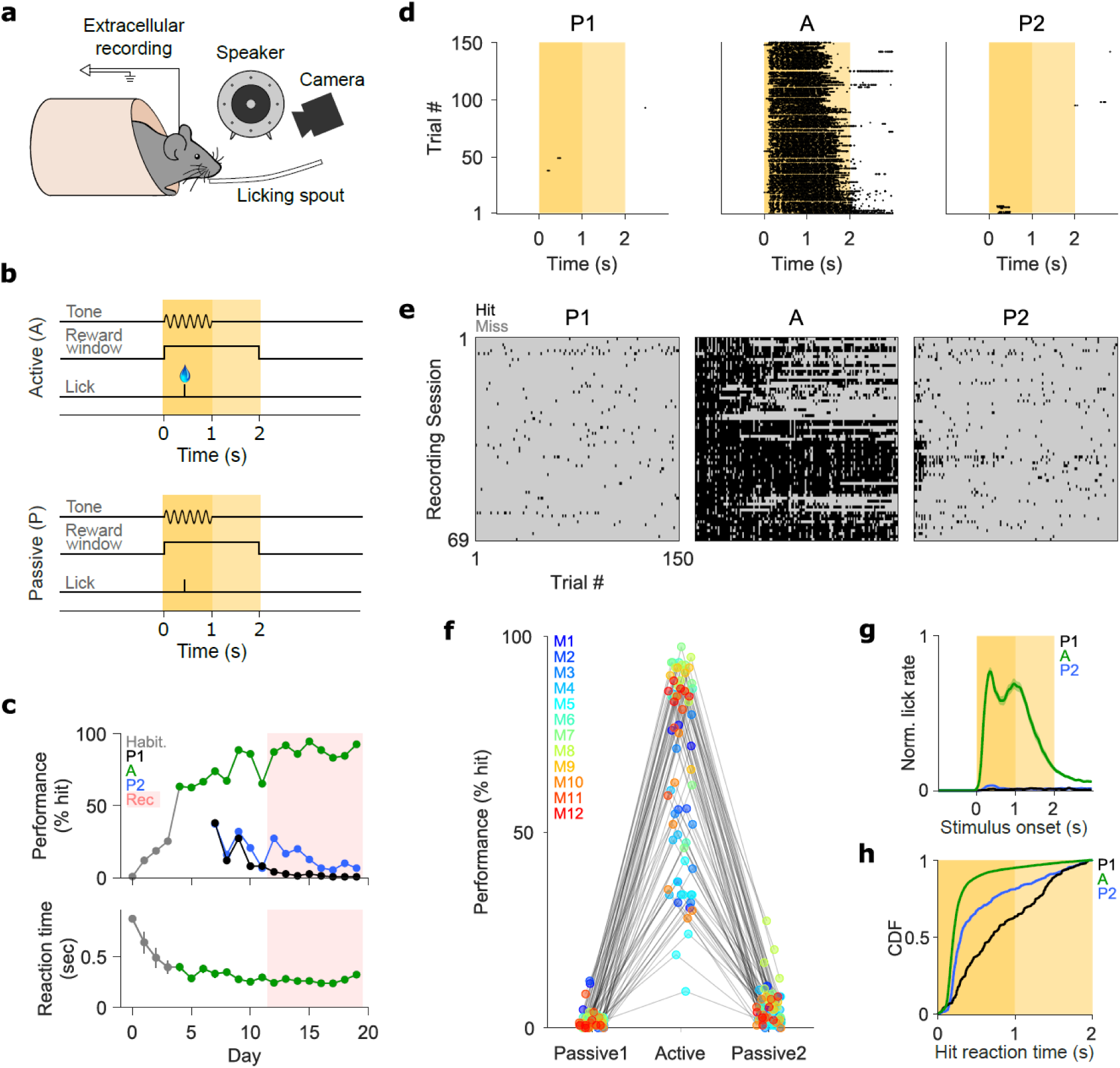
Mouse performance is reliably different between passive and active phases. **a**, Schematic of the experimental setup for recording in awake, behaving animals. Mice were head-fixed in a carton tube in front of a licking spout delivering the reward. A loudspeaker was positioned 10 cm away from the left ear and delivered tones. Pupil size was imaged using a high-definition camera next to the speaker. Multi-channel extracellular electrophysiological recordings were performed in the right hemisphere. **b**, Schematic of the behavioral paradigm. Each session consisted of one active phase flanked by two passive phases. In the active phase, mice had to lick within a 2 s reward window (yellow shaded area) in response to 1 s pure tones (dark yellow shaded area) of varying frequency to get a reward. In the passive phase, no reward was released. The inter-trial interval varied between 3-7 s, and the timer was reset if the animal licked the reward spout outside of the reward window. **c**, Learning curve for a representative mouse across training (white area) and recording (red shaded area) days. Grey points: habituation phase. Green: active blocks (A). Black: first passive block (P1, preceding active). Blue: second passive block (P2, following active). Top: performance was measured as the percentage of correct trials over the total trials in each block. Bottom: average reaction time for hit trials during the active phase. **d**, Example of one session from one mouse. Each row represents one trial in P1 (left), A (middle) or P2 (right) blocks. Each dot represents when the piezo signal exceeded the detection threshold tracking licking behavior. **e**, Hit (black) and miss (grey) trials for all recording sessions in P1 (left) A (middle) and P2 (right) phases. Each row depicts one recording session in one mouse. **f**, Performance in P1, A, and P2 phases. Each color represents one mouse. Each triplet of connected dots represents performance in one of the 69 recording sessions. **g**, Average normalized licking behavior during P1 (black), A (green), and P2 (blue). Licking behavior was averaged across trials in each phase and session, normalized across phases in each session, and then normalized across sessions. **h**, Cumulative distribution function of reaction times in hit trials during P1 (black, n = 184), A (green, n = 6757) and P2 (blue, n = 506) phases. See also Figure S1.

### Neural responses to pure tones are robust along the auditory pathway

We investigated how task-engagement influences auditory neurons using single or multi-shank, 32 or 64 channel silicon probes to perform acute extracellular recordings in awake mice executing the tone-detection task described above. We recorded 4414 single and multi-units from IC (n = 622), MGB (n = 654), ACx (AAF, n = 1439; A1, n = 1145) or unidentified nearby regions (n = 554) (Figure 2a, d) of 12 mice in 69 behavioral sessions (Table S1). Cortical neurons were classified as belonging to AAF or A1 based on tonotopy, their laminar position was identified as superficial (Sup), input (Inp) or deep (Deep) based on current source density analysis (Figure 2b), and their neural subpopulation as putative regular (RS) or fast-spiking (FS) based on the peak-to-trough duration of their action potential (Figure 2c) (see Methods). This yielded recordings from 14 anatomically identified groups of neurons (Figure 2d).

**Figure 2.**
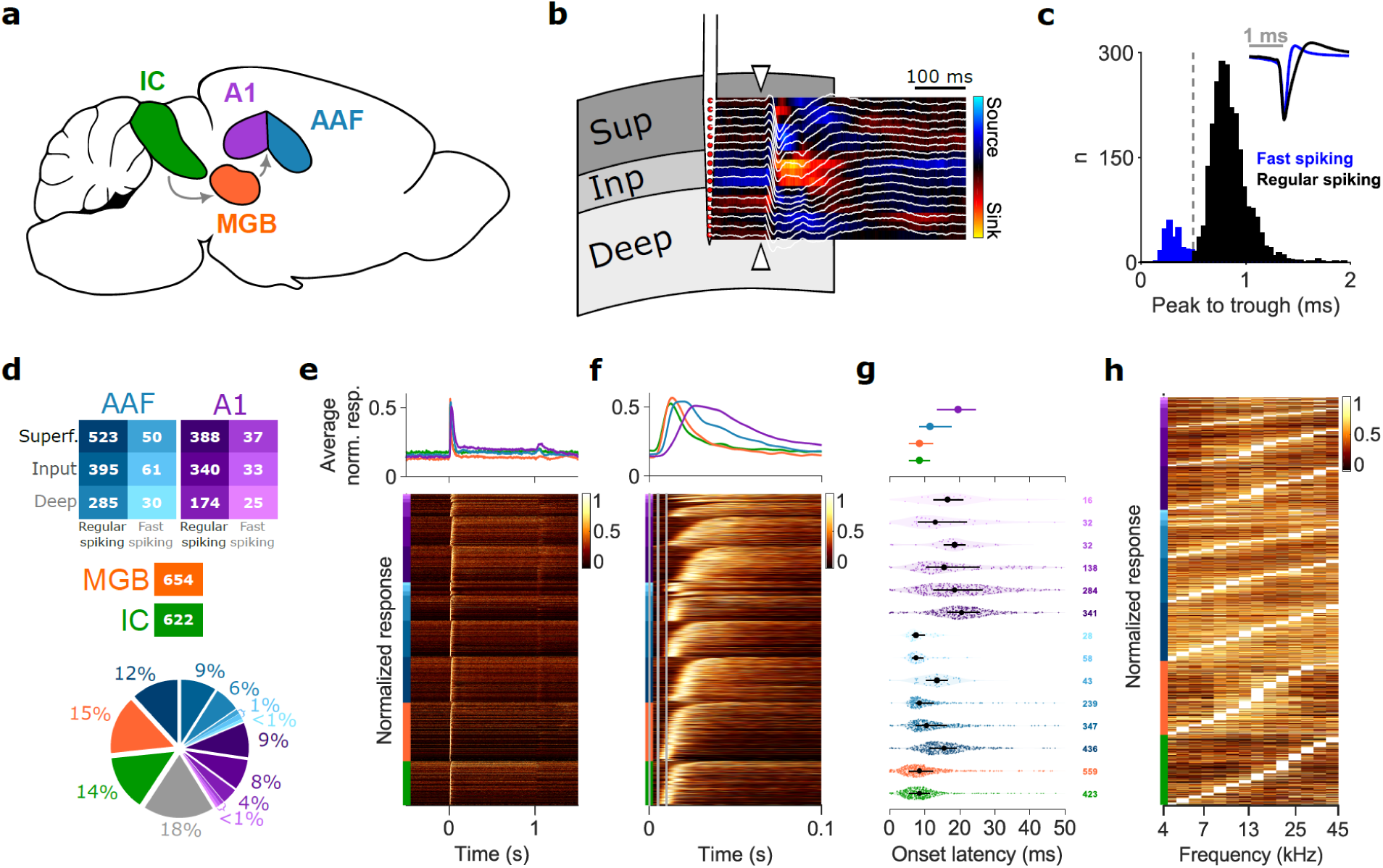
Neural responses to pure tones are robust along the auditory pathway. **a**, Schematic of the regions from which neurons were recorded: IC (green), MGB (orange), A1 (purple), or AAF (blue). **b**, Laminar localization of recording sites in primary auditory cortices (A1 and AAF) via current source density analysis (red-blue map) of local field potentials (LFP, white traces) in response to 50 ms long white-noise bursts (onset indicated by white arrows). Channels were classified as located in input layers if the relative LFP was part of the largest sink (warm colors), and as superficial or deep layers otherwise, depending on the relative position to sink channels. **c**, Histogram of action potentials’ peak-to-trough duration in cortical putative fast-spiking (blue, n = 261) and regular-spiking (black, n = 2323) units. The dotted line indicates the threshold for classification (0.5 ms). Inset: average spike shape across fast-spiking (blue) and regular-spiking (black) units. **d**, Top: schematic depicting color code used for anatomical identity across figures, and numbers of units for each group. Bottom: proportion of recorded units for each group. Grey: units recorded in unidentified cortical regions. **e, f**, Responses to 1s long pure tones in the first passive phase (P1) around tone onset (0 s) (e) and zoomed version around onset responses (first 100 ms following tone onset) (f). Top: average peristimulus time histogram (PSTH) across units recorded in each area (color code as in (d)). Responses were averaged across all trials for each unit, normalized, and then averaged across neurons in each area. Bottom: normalized average PSTHs across trials for each anatomically identified unit (n = 3617). Each row represents one unit. Units are ordered from bottom to top by anatomical identity (color bars on the left, color code as in (d)), and by response onset latency. In (f), vertical grey lines indicate 0, 5 and 10 ms. **g**, Onset latency of tone-evoked responses (color code as in (d)), for units in which onset response could be estimated (see Methods). Dots indicate medians, lines indicate 25^th^ and 75^th^ percentiles. Top: Distribution of onset latency in each area. Bottom: violin plots of onset latencies in each anatomically identified group. Each colored dot is one unit. The number of units per group is indicated on the right. **h**, Normalized frequency tuning for each anatomically identified unit (n = 3617) during the first passive phase (P1). For each unit, responses were averaged across trials of identical tone frequency and then normalized across frequencies. Units are ordered from bottom to top by anatomical identity (color bars on the left, color code as in (d)), and by frequency eliciting maximal response.

Responses to pure tones were robust across the auditory pathway and characterized by a vigorous onset response that greatly dampened within the first 100 ms (Figure 2e-f). As expected, the response latency increased along the pathway (Figure 2f-g). The frequency selectivity of the recorded units spanned the whole range of tested frequencies in each area (Figure 2h), confirming appropriate sampling across regions.

### Neural responses across the auditory pathway are modulated by task-engagement

We first asked if engaging in the tone-detection task modulates neuronal activity. To investigate this, we compared the activity of individual units around tone onset in the active and passive phases of the task. We found a great variety of response modulations (Figure 3a) including changes in the onset auditory response that could be accompanied by frequency tuning adjustments, and changes in the sustained responses that could be precisely timed to the tone duration or persist after tone offset.

**Figure 3.**
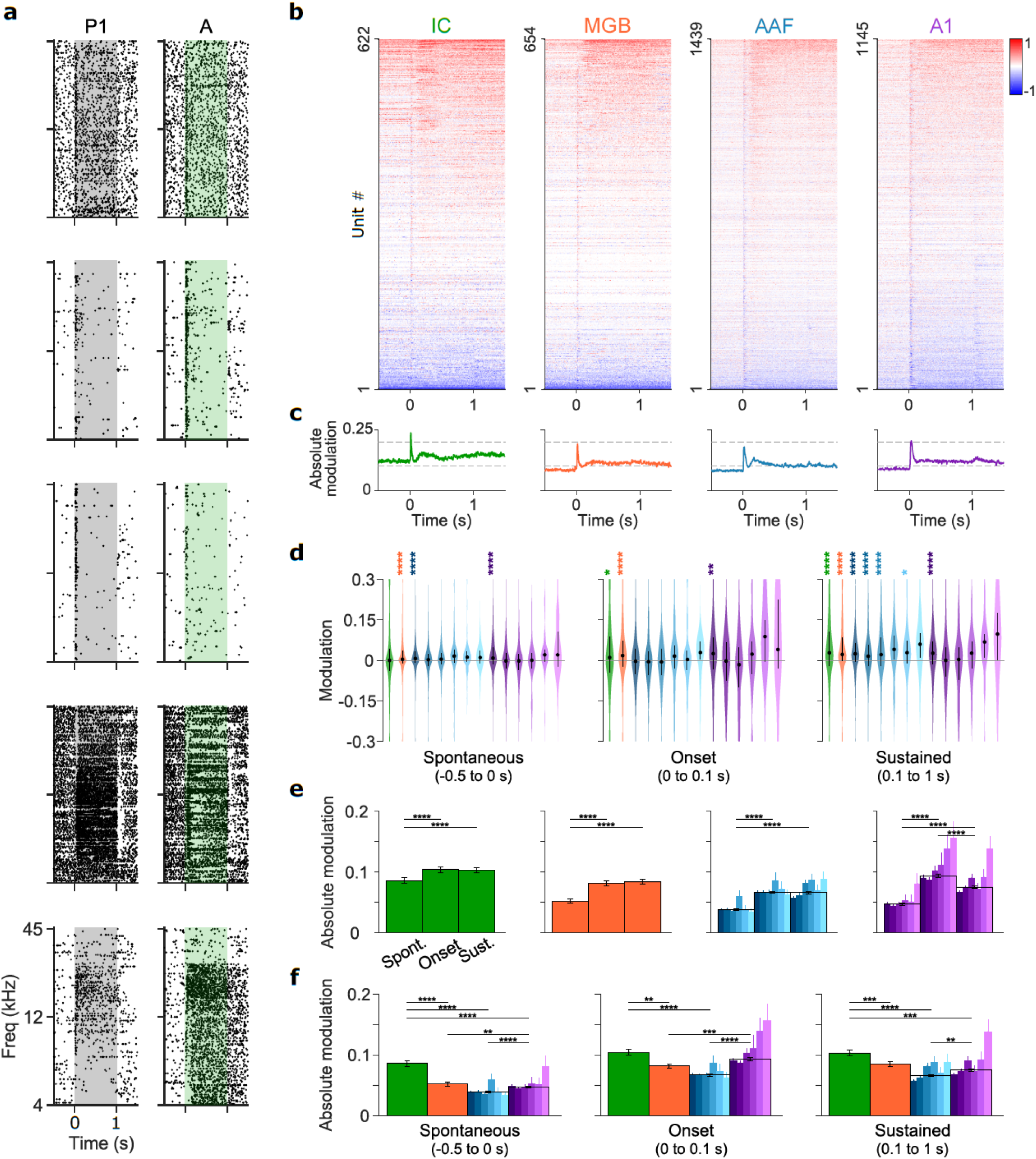
Neurons across the auditory pathway are modulated by task engagement. **a**, Raster plots showing the activity of 5 example units around tones of different frequency in the first passive (left) and active phase (right). Shaded areas indicate periods of tone presentation. Each dot represents one spike, each row represents one trial. Trials are ordered by tone frequency. **b**, Modulation PSTHs depicting the change in activity around tones from P1 to Ahit in each area. In each area, each row is the modulation PSTH of one recorded unit. Red indicates an increase in activity in the active phase, blue indicates a decrease in activity. **c**, Population average of absolute modulation PSTH in each area. Dashed lines indicate modulations of 0.1 and 0.2. **d**, Violin plots showing distributions of average task-induced modulations (calculated for each unit from modulation PSTH shown in (b)) of spontaneous rate (−0.5 to 0 s around stimulus onset, left), onset response (0 to 0.1 s, middle) and sustained responses (0.1 to 1 s, right) in each anatomically identified group. Color code as in Figure 2d. Black dots and lines indicate median, 25^th^ and 75^th^ percentiles, respectively. Asterisks indicate significant differences from 0 for each anatomically identified group (Wilcoxon signed-rank test, *m*=42). **e**, Histograms of absolute modulation of spontaneous (left bar), onset (middle bar) and sustained (right bar) rates in each area (IC, green; MGB, orange; AAF, blue; A1, purple). For AAF and A1, the average is depicted by black outlines and layer/neuronal subclasses break down is depicted by individual colored bars (color code as in Figure 2d). Data represent mean ± SEM. Wilcoxon signed-rank test, *m*=12. **f**, Same data as in (e), but highlighting the comparison of spontaneous, onset and sustained modulations between different areas. Wilcoxon rank-sum test, *m*=18. See also Figure S2.

We then asked how the modulations we observed varied along the auditory pathway. Since we expected the largest modulations to happen between the first passive (P1) and the correct trials of the active phase (Ahit), we initially focused on the changes in responses between P1 and Ahit trials. To do so, we calculated the difference between the normalized average activity in P1 and Ahit trials (Figure S2) (see Methods). We found widely distributed engagement-modulations in all brain regions (Figure 3b). Such changes were already visible in the spontaneous activity preceding tone onset, but they became particularly evident in the form of strong and fast modulations of tone onset responses and also as slower, more sustained changes later in time (Figure 3b-c). We therefore subdivided the temporal profile of task-engagement modulations into three periods: prestimulus (−0.5 to 0 s before tone onset), early (0 to 0.1 s) and late (0.1 to 1 s) to distinguish the average changes in spontaneous rate, onset or sustained responses, respectively. The changes in activity ranged widely and varied from suppression to enhancement in all areas, while sustained modulations tended to be of positive polarity (Figure 3d). The magnitude of modulation was significantly stronger after tone onset, and it was similar between onset and sustained responses in all areas apart from A1, where onset modulations were the largest (Figure 3e). Surprisingly, task-engagement modulations were generally more vigorous in IC than in downstream areas (Figure 3f). AAF consistently displayed the lowest level of modulation. Interestingly, the largest A1 onset and sustained modulations happened in deep layers (Figure 3e-f).

In summary, we found widespread and diversified task-induced modulations across the auditory pathway that were stronger after sound onset.

### Task-engagement induces distinct modulation motifs in different clusters of neurons

The temporal partitioning of changes in spontaneous, onset and sustained activity provides a coarse description of how task-engagement modulates neurons along the auditory pathway, but is also likely to hide more temporally precise modulations. For example, late modulations can have different temporal dynamics: in some units they are restricted to the stimulus period, in others they persist even after tone offset (Figure 3a), while in others they result in a bump of activity slowly ramping up and then decaying within a few hundred milliseconds (see IC and AAF, Figure 3b-c). To identify the different types of changes in activity induced by task-engagement, we performed clustering analysis of such modulations (see Methods). We found ten clusters displaying distinct motifs of functional modulation and aggregated the neurons that were not reliably grouped with any other neuron into an eleventh cluster (Figure 4a-d, Figure S3). Most neurons belonged to one of the clusters that displayed consistent engagement modulations (3692/4414 units, 83.6%). The clustering algorithm segregated the neural modulations mainly based on their temporal dynamics (onset-sustained) and their polarity (enhancing-suppressive) (Figure 4c-d, Figure S3).

**Figure 4.**
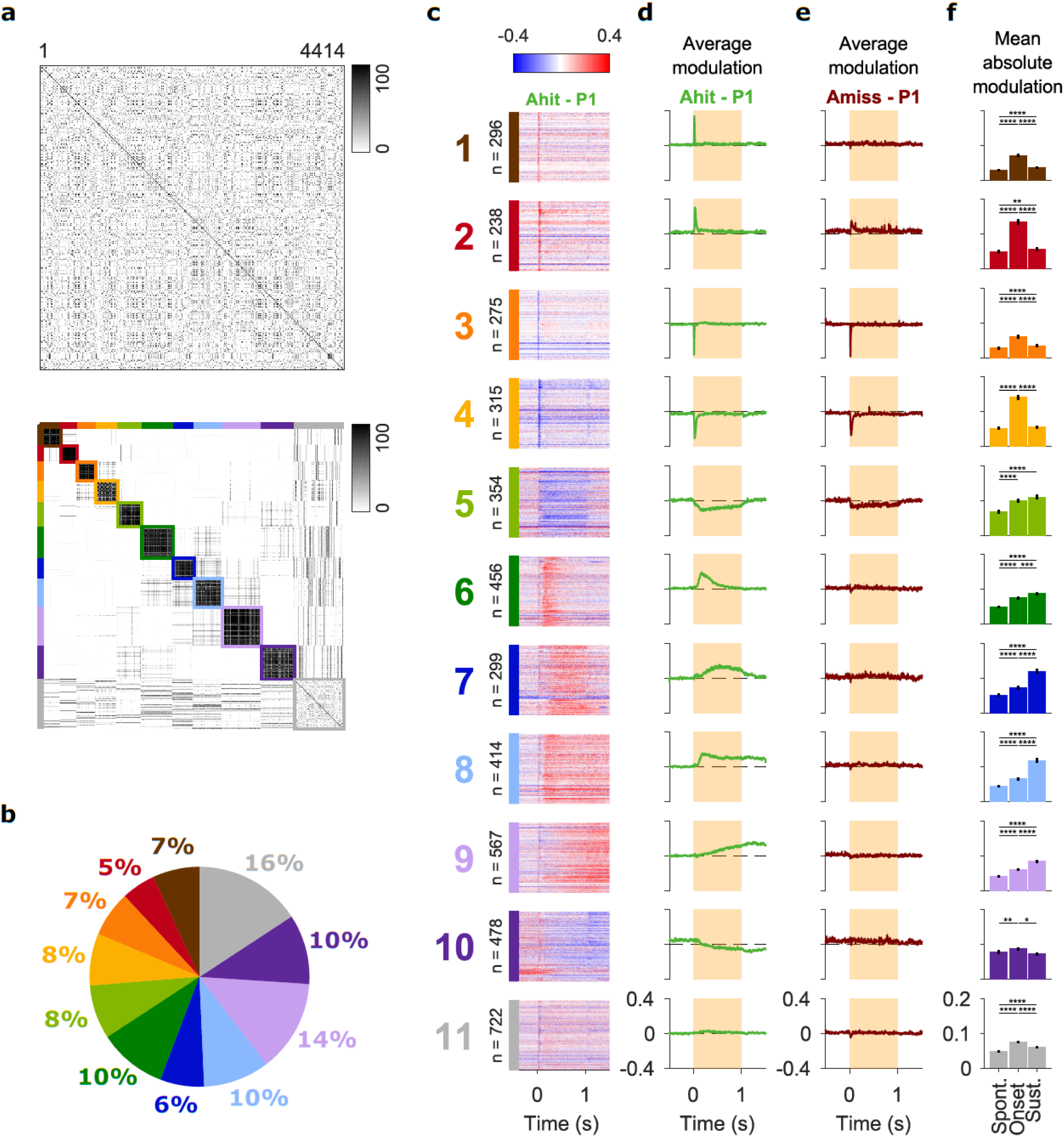
Task-engagement induces distinct modulation motifs in different clusters of neurons. **a**, Top: co-clustering matrix resulting from reiterated K-means clustering analysis of response modulations for all recorded neurons (n = 4414). The color represents the percentage of K-means runs in which each neuron was assigned to the same cluster of another neuron (see Methods). The order is based on recording sessions. Bottom: result of hierarchical clustering of co-clustering matrix data shown in (Top). Neurons are sorted based on the cluster they were assigned to. The identified final clusters are highlighted by colored squares and lateral bars. **b**, Percentage of recorded neurons assigned to each functional cluster. **c**, Modulation PSTH of all neurons assigned to each functional cluster (color code as in (a, b)), one neuron per row. The numbers on the left indicate the number of neurons assigned to each cluster. **d-e**, Average modulation PSTH from trials in the first passive phase to correct hit (d) or miss (e) trials in the active phase for neurons belonging to each cluster. Shaded areas indicate tone duration. Dashed horizontal grey lines indicate 0 modulation. **f**, Absolute spontaneous, onset and sustained average modulation across units in each cluster. Wilcoxon signed-rank test, *m*=33. Data show mean ± SEM. See also Figures S3 and S4.

All clusters displayed robust onset response modulations (Figure 4d, f and Figure S3). Some clusters predominantly displayed such onset modulations (#1-4), and were further subdivided in very brisk (#1, 3) or slower (#2, 4) decays. These primarily onset modulations were found in about one-fourth of the recorded neurons (1124/4414, 25.5%). Neurons in other clusters also displayed vigorous modulations of the onset response, but they were clustered separately due to concurrent late modulations. Since early and late modulations in these clusters did not always have the same polarity, onset modulations were washed out in the average modulation peristimulus time histograms (PSTHs) (Figure 4d, Figure S3). Onset modulations corresponded to the period in which the largest auditory responses were found (Figure 2e-f, Figure S3a) while motor actions, and therefore reward, still had to happen (Figure 1c, h), making them the likely substrate of the perceptual changes arising during listening. Neurons belonging to cluster 5 also displayed late modulations, but they had short onset latency and were precisely restricted to the stimulus period. Since they occurred in neurons characterized by robust auditory responses that lasted throughout the tone presentation rather than being transient (Figure S3a), they are likely also influencing sound processing.

The other clusters were characterized by additional changes occurring in the late phase of the response (#6-10) and included 50.2% of the recorded neurons (2214/4414). These changes could arise quickly after tone onset (#6, 8, 10) or have a slower progression (#7, 9), and they were further subdivided into decaying (#6-7) or persistent (#8-10) modulations. These task-induced changes in neural activity happened in periods where passive auditory responses were much lower than in the first 100 ms (Figure 1d, Figure S3a). The timing of these modulations overlapped with the animals licking the spout and receiving a reward, suggesting that they may reflect the integration of motor or reward signals in the auditory system.

### A subset of modulation patterns correlates with behavioral performance

We next asked which modulations may help auditory perception. To do so, we compared the observed changes in hit and miss trials (Figure 4d-e). We found a clear difference between early and late modulations: late ones were completely absent in miss trials (#6-10), supporting the hypothesis that they reflected either licking or the presence of reward. Intriguingly, the amplitude of pure onset changes was more variable: enhancements of auditory responses were either gone or heavily weakened in miss trials (#1-2), while suppressive modulations were relatively similar between correct and miss trials (#3-5). This finding was also true when we considered the whole population of neurons (Figure S4b). The selectivity of onset response enhancement for correct trials suggests they may play an important role in improving auditory perception during active listening. The preservation of suppressive modulations in both hit and miss trials suggests instead that they may be less important for auditory perception, but rather reflect a brain-state change from passive to active hearing that does not influence hearing performance.

Previous studies have shown that task-induced plasticity in the auditory system often persists even after the animal stops performing a task (Atiani et al.,2009; Fritz et al., 2003; Fritz et al., 2005, 2007; Slee and David, 2015). To investigate the degree of persistence of the observed modulations, we compared the level of modulation between the first and the second passive phases (Figure S4a, c). Consistent with previous results, modulations of the auditory onset responses generally persisted after the animals stopped licking in response to the tones. Nevertheless, the amplitude of such modulation was decreased, confirming that the modulations depend on the level of task-engagement. As observed for miss trials, late modulations were absent from the second passive phase.

### Task-engagement can modulate gain control and frequency tuning

To study how task-engagement modulates the excitability and the sound sensitivity of auditory neurons, we investigated their changes in gain (Ferguson and Cardin, 2020; Phillips and Hasenstaub, 2016) and frequency tuning. To do so, we selected the units in which the tuning curve of tone onset responses could be approximated by a Gaussian fit in both P1 and Ahit (1462/4414) (Figure 5a).

**Figure 5.**
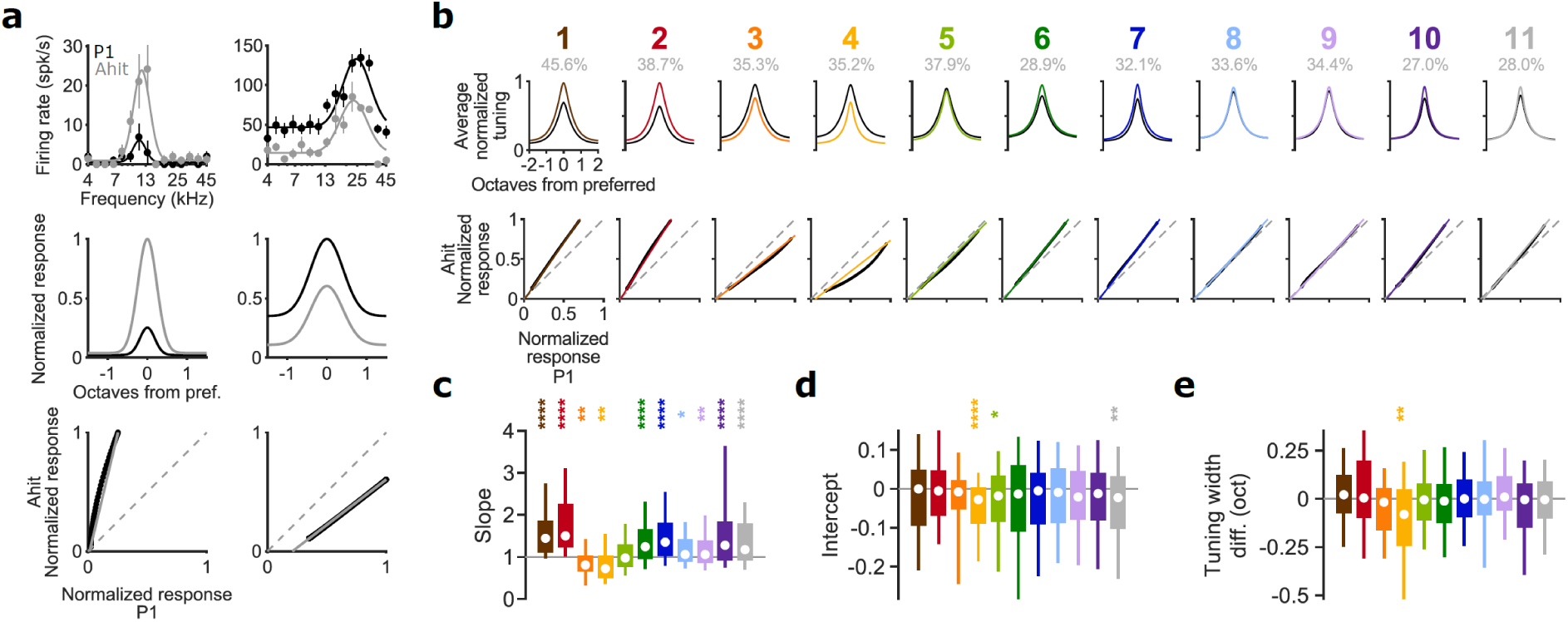
Task-engagement can affect gain control and frequency tuning. **a**, Pipeline of frequency tuning and gain change estimation for two example neurons. Top: Average response to tones at each frequency in the first passive (black, P1) and hit trials of active phase (grey, Ahit) (dots and lines are mean ± SEM, respectively). Solid lines depict the best-fit Gaussian model. Middle: best fit Gaussians were aligned to the preferred frequency and normalized between P1 and Ahit. Bottom: normalized, centered fitted responses of P1 fit are plotted against those of Ahit (black points). Grey solid line depicts the linear fit of the lowest and highest responses. The dashed grey line is the equality line (no gain changes). The unit on the left shows pure multiplicative gain changes. The unit on the right shows a mixture of divisive and subtractive gain changes. **b**, Top: average tuning curves in first passive (black) and hit trials of the active phase (colored curves) for each cluster (color code as in Figure 4), after normalizing between P1 and Ahit in each unit and centering to preferred frequency. Grey percentage indicates amount of units with good Gaussian fit. Bottom: linear fit (colored line) of average P1 against Ahit tuning curves (black) shown in (top). **c**,**d**,**e**, Box plots showing distribution of estimated (c) slopes (multiplicative/divisive gain changes), (d) intercepts (additive/subtractive gain changes) and (e) tuning width modulations (broadening/sharpening of tuning curve) in each functional cluster. Color code as in Figure 4 and (b). In each boxplot, white dot indicates median, rectangle 25^th^ and 75^th^ percentiles, whiskers 10^th^ and 90^th^ percentiles. Dashed grey horizontal lines indicate no change. Wilcoxon signed-rank test, *m*=33. See also Figure S5.

We estimated multiplicative and additive gain changes as the slope and intercept, respectively, of the linear fit to maximum and minimum firing rates of normalized tuning curves aligned to peak in the two behavioral states (Figure 5a). In most clusters, we found significant multiplicative gain changes (all except #5). The largest multiplicative gain changes were found in clusters that displayed enhanced onset responses (#1-2): these were the modulations that correlated with behavioral performance (Figure 4d-e and Figure S4b), suggesting that the neural mechanisms underlying multiplicative gain changes may be important for auditory perception during active listening. Neurons whose onset activity was suppressed in the engaged state displayed divisive gain modulations instead (#3-4) (Figure 5b-c). Task-engagement also induced moderate subtractive tuning shifts in some clusters (#4, 5, 11) (Figure 5b, d). Gain and spontaneous activity modulations correlated in many clusters (Figure S5), suggesting that gain changes are likely to be at least partially due to general changes in neural excitability. To estimate changes in frequency selectivity that cannot be captured by the gain changes described above, we analyzed changes in tuning width (estimated as the Gaussian fits’ sigma) (Figure 5e). Interestingly, only those clusters with onset response suppression and with divisive gain modulations (#4) consistently displayed tuning width changes resulting in sharpened tuning curves in the active phase.

In summary, our results suggest that enhancing and suppressive onset modulations result from distinct sets of gain changes, and will help formulate and test hypotheses to dissect the underlying neural mechanisms.

### The emergence of onset auditory modulations follows the canonical auditory pathway

Modulations of the onset auditory responses were found across the auditory pathway (Figure 3b-f). However, changes in subcortical response amplitudes may arise from the massive feedback from primary ACx. If they were, they would appear after cortical modulations. To test this, we analyzed the latency of onset modulations. Inconsistent with the hypothesis of cortical feedback, onset modulations in subcortical regions emerged before cortical ones (Figure 6a). The emergence of early task-engagement modulations therefore follows the canonical auditory pathway, implying that they do not arise in primary auditory cortices to then be fed back to subcortical regions. Instead, cortical onset modulations could even just be inherited from subcortical stations.

**Figure 6.**
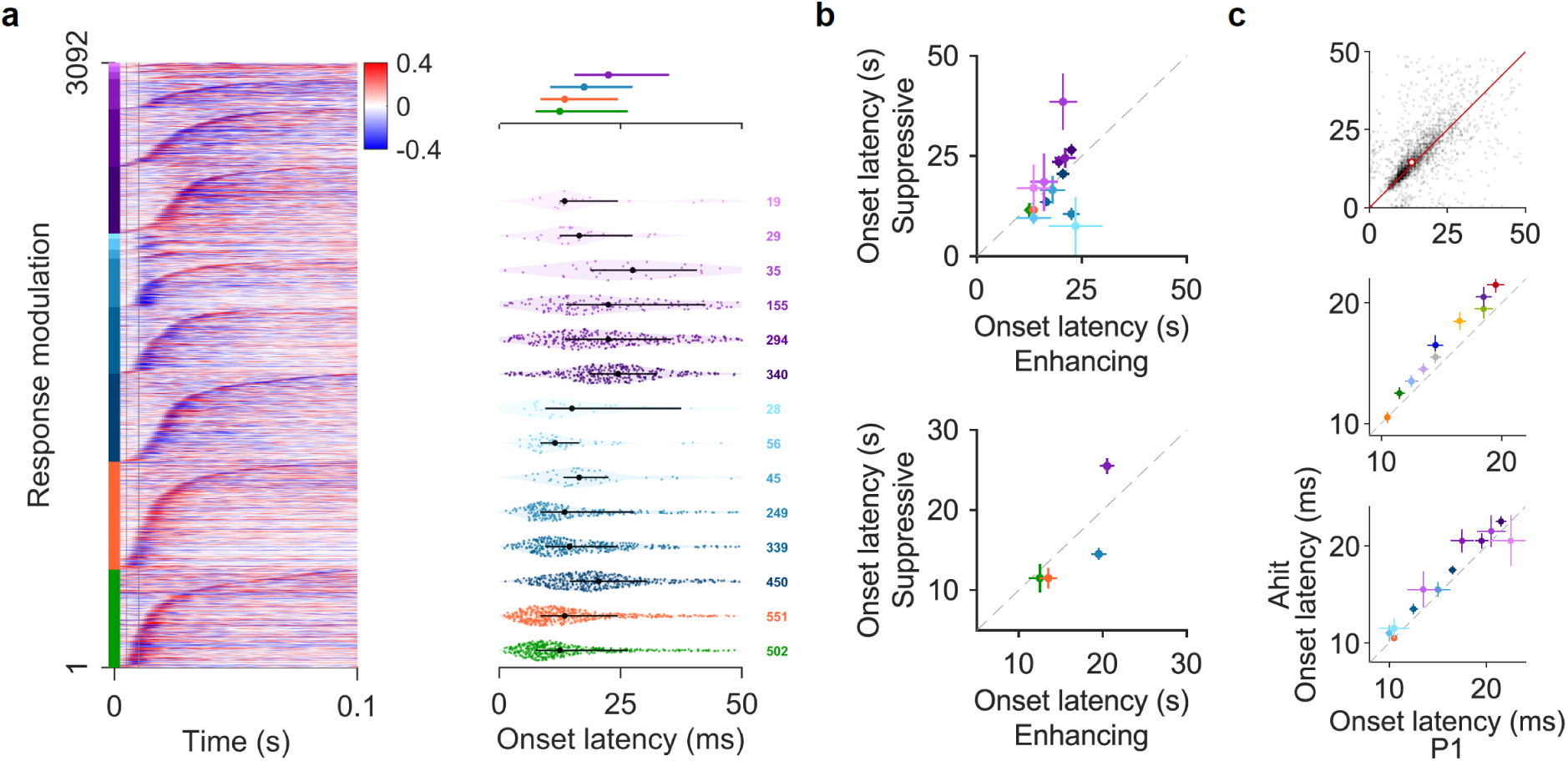
The emergence of onset auditory modulations follows the canonical auditory pathway. **a**, Onset latency of task-induced modulation along the auditory pathway. Left: onset modulation PSTHs for each anatomically identified unit in which onset of modulation could be measured (n = 3092, see Methods). Each row represents one unit. Units are ordered from bottom to top by anatomical identity (color bars on the left, color code as in (Figure 2d), and by modulation onset latency. Vertical grey lines indicate 0, 5 and 10 ms. Right: distributions of modulation onset latency in anatomically identified regions. Dots indicate medians, lines indicate 25^th^ and 75^th^ percentiles. Top: Distribution of onset latency in each area. Bottom: violin plots of onset latencies in each anatomically identified group. Each colored dot is one unit. The number of units per group is indicated on the right. **b**, Latency of enhancing against suppressive onset modulations in each anatomically identified region (top) or area (bottom). Wilcoxon rank-sum test, *m*=16 (See Table S2). **c**, Scatter plots of latency of onset (0-100ms) tone-evoked responses in first passive phase (P1) against hit trials in active phase (Ahit) for units in which it could be measured in both (see Methods). Top: all anatomically identified units (n = 2303). The red dot depicts median ± SEM. Latency was larger in Ahit trials (p < 10^−21^, Wilcoxon signed-rank test). Middle: onset latency for each functional cluster (n = 2683). Wilcoxon signed-rank test, *m*=11 (see Table S3). Bottom: onset latency for each anatomically identified area. Wilcoxon signed-rank test, *m*=14 (see Table S4). Data show median ± SEM in (b-c).

We next asked if enhancing and suppressive onset modulations followed a precise temporal sequence. We found that suppressive modulations were consistently faster than enhancing ones in AAF, while the contrary was true in A1. Subcortical regions instead showed no difference in latency (Figure 6b) (Table S2).

We also observed that tone response latency increased in the active phase across the population of neurons we recorded (Figure 6c). The latency increment was found in most clusters (#3-4, 6-8, 10-11) (Table S3), and in most cortical regions (Table S4), but again it was absent in subcortical areas. These results suggest that some of the neural mechanisms responsible for task-induced response modulations are selectively implemented in cortical circuits.

### Late modulations can be induced by arousal, movement or reward

Onset response modulations happened while mice were attentive but not yet licking in response to the tones (Figure 1h), and therefore reflected task-induced changes mainly acting on the strongest auditory responses (Figure 2e-f, Figure S3a). In contrast, late response changes occurred when mice were also licking the spout and receiving a reward, both of which likely correlate with a state of increased arousal. To disentangle the origin of such late modulations, we extracted epochs characterized by nested behavioral states from the active phase (see Methods). We first selected epochs aligned to the first lick of hit trials (−1 to 3 s around the first lick), characterized by prolonged licking behavior and the presence of reward. As expected, a lick accompanied by reward induced a surge of arousal revealed by an increase in pupil radius (Figure 7a-c). Clusters with stronger onset modulations (#1-5) displayed the largest peak before lick onset (Figure 7f), reflecting the responses to the tones blurred as a consequence of the variable hit latency. Interestingly, the activity of clusters characterized by late modulations (#6-10) peaked at or after lick onset. The activity of cluster 6, culminating at lick onset and then decaying, is consistent with the hypothesis that neurons belonging to this cluster may integrate motor-preparation or reward-prediction signals. In contrast, other late-modulated clusters (#7-10) showed more sustained responses, suggesting that they were sensitive to either licking, reward or arousal (Figure 7f).

**Figure 7.**
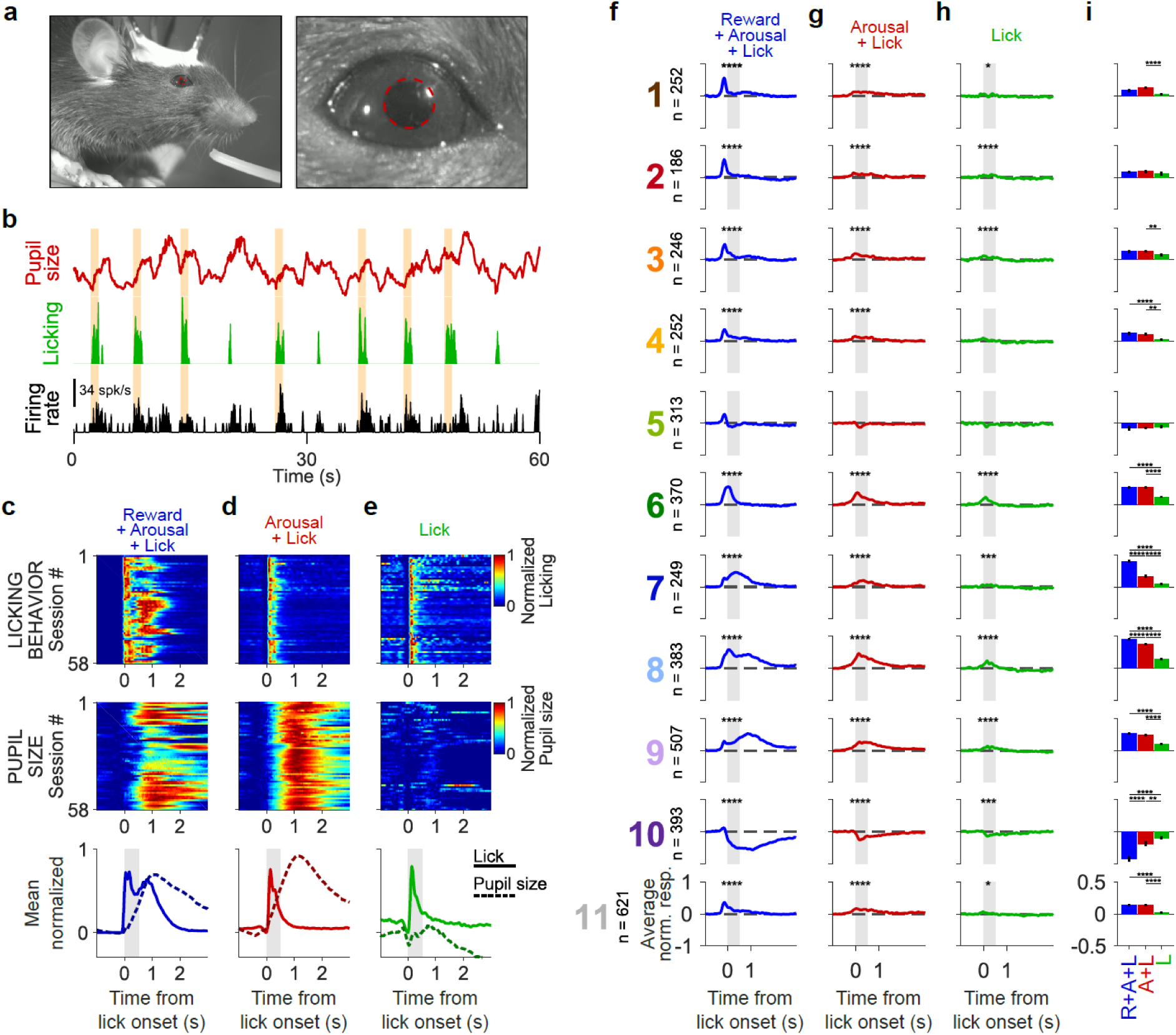
Late modulations can be induced by arousal, movement or reward. **a**, Pupil size was monitored by a high-definition camera and extracted off-line with DeepLabCut (Nath et al., 2019). Left: Example frame from video monitoring of one mouse during behavior. Right: Zoomed in version on the mouse eye. The red dashed line shows the estimated pupil size and position. **b**, Example traces of estimated pupil size (top, red), licking behavior (middle, green) and PSTH of one unit (bottom, black) for 1 minute recording in the active phase. Yellow dashed vertical areas indicate tones. **c, d, e**, We extracted recording epochs (−1 to 3 s) around the first lick in hit trials of the active phase (a), or around the onset of spontaneous licking bouts away from tone presentation accompanied by arousal increases (d) or not (e). Top: average normalized licking behavior in each recording session. Middle: average normalized pupil size in each recording session. Bottom: average normalized licking behavior (solid lines) and pupil size (dashed lines) across sessions in different epochs. Shaded grey area indicates the window (0-400 ms) used for the analysis of neural responses. **f, g, h**, Average normalized PSTH in epochs shown in (c,d,e) for each functional cluster. Each row corresponds to one cluster (color code as in Figure 4). The number of units indicated on the right of cluster number. Dashed grey area indicates the window (0-400 ms) used for the analysis of neural responses in each epoch (see also panel (i)). Asterisks indicate significant responses. Wilcoxon signed-rank test, *m*=33. **i**, Comparison of mean ± SEM neural responses in the different behavioral epochs shown in (c,d,e) for each cluster. Wilcoxon signed-rank test, *m*=33. Data in (c-e, bottom) and (f-i) show mean ± SEM. See also Figure S6.

To uncouple reward and tone responses from movement and arousal signals, we next extracted epochs around licking bouts that happened far from the tones during the active phase (Figure 7b, d-e, g-h). To dissociate modulations caused by changes in arousal from those caused by licking behavior, we further subdivided these licking epochs based on the concomitant changes in pupil size. This resulted in a second behavioral state in which licking that occurred in the absence of tones and reward was accompanied by an increase in pupil radius (Figure 7d,g) and in a third behavioral state where there was no clear change in pupil size (Figure 7e,h). Analyzing average neural responses (0.4 s following lick onset) revealed that most clusters were moderately but consistently modulated by licking (#1-3, 6-11) (Figure 7g-i). Comparing lick responses in the presence and absence of pupil changes revealed that arousal also influenced many clusters (#1, 3-4, 6-11) but not all of them (#2, 5) (Figure 7g-i). In contrast, the presence of reward only affected a subset of clusters (#7, 8, 10) (Figure 7f, g, i). In summary, we found that the different patterns of late modulations we observed in hit trials in the active phase of the task could be explained by cluster-specific levels of neural sensitivity to licking behavior, arousal or reward.

### Task-engagement modulations are distributed along the auditory pathway

The task-induced modulations may start to appear only at a certain stage along the auditory pathway, or be increasingly represented at higher stages of sensory processing. If specific modulations were mainly to be found in particular brain regions, it would help us elucidate the role these regions play in different aspects of task-engagement. To answer this, we investigated the anatomical distribution of task-induced modulations. To our surprise, neurons belonging to each functional clusters we observed were found across the auditory nuclei that we investigated (Figure 8a). This finding shows that all the task-induced modulations found in cortex are also found in subcortical auditory stations as early as the IC.

**Figure 8.**
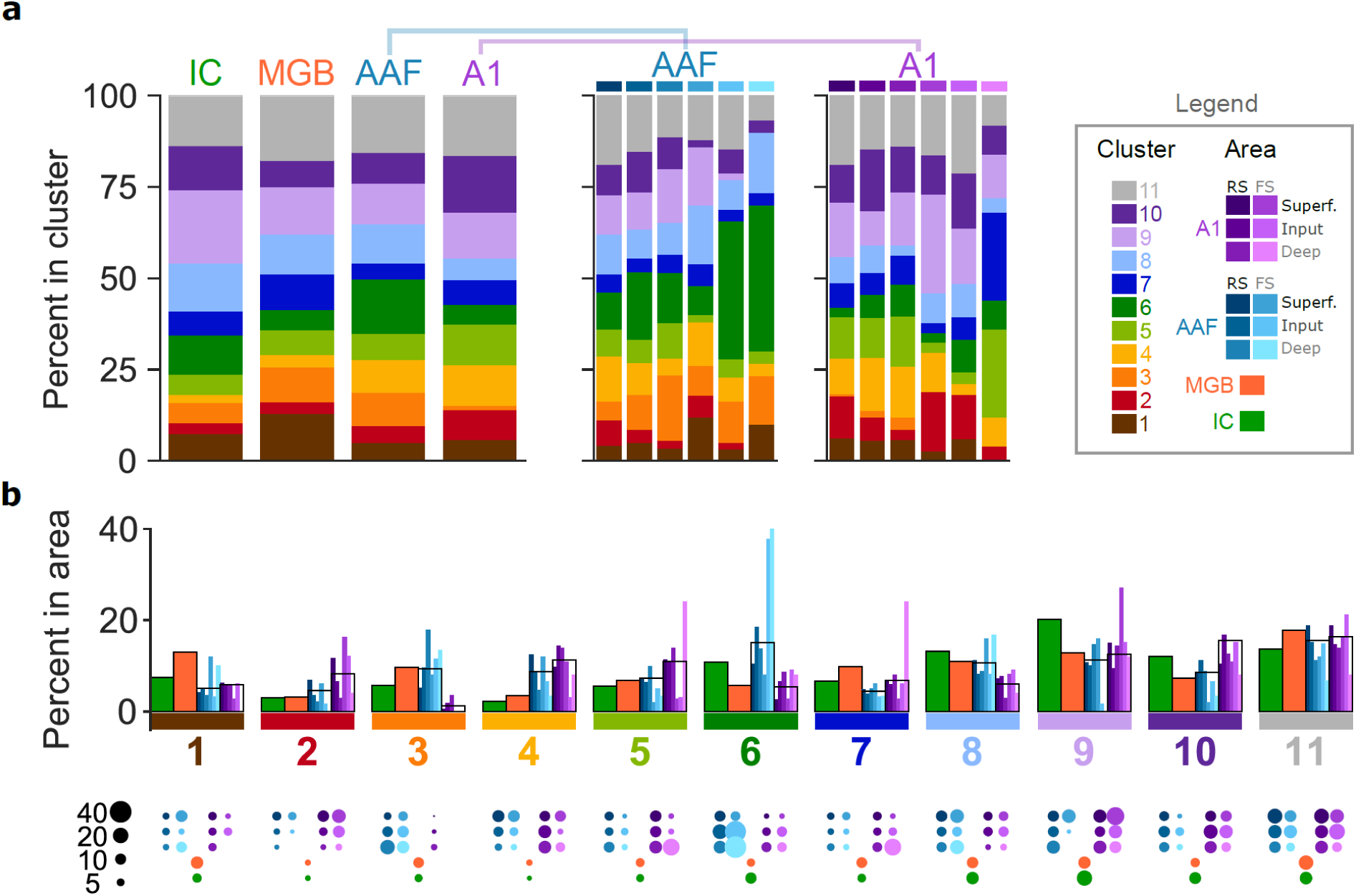
Task-engagement modulations are distributed along the auditory pathway. Color code for anatomical groups as in Figure 2d, for functional clusters as in Figure 4. **a**, Left: percentage of neurons belonging to each functional cluster in each area (IC, n=622; MGB, n=654; AAF, n=1439; A1, n=1145). Middle, right: Percentage of neurons belonging to each functional cluster in different layers and neuronal subclass of AAF (middle) and A1 (right). **b**, Top: Percentage of neurons belonging to each functional cluster in each area. Same data as in (a), displayed by cluster rather than by area. Bottom: bubble plot displaying the percentage of neurons belonging to each functional cluster in each area (same data as (a,b top)). The larger the bubble, the larger the percentage (scale bar on the left). See also Figure S6.

A more fine-grained analysis revealed distinct aspects of the anatomical segregation of functional clusters (Figure 8b). Some clusters were increasingly represented along the auditory pathway (#2, 4, 5), while others were decreasingly represented and mainly found in IC (#8-9). Other clusters were particularly abundant in fast-spiking neurons in input and deep layers of AAF (#6) or fast-spiking neurons in deep layers of A1 (#5, 7). Clusters 2 and 4 were decreasingly represented from superficial to deep cortical layers. Clusters 3, 6 and 8 were mainly found in AAF rather than A1, while the opposite was true for clusters 2, 4, 5 and 10. Consistent with increasingly longer dynamics of onset responses to sound along the auditory pathway (Figure 2e-g), we found that the slower onset modulations (#2, 4, Figure 4c-d) were also progressively more represented.

Neurons showing the largest responses to licking behavior (#6, 8) (Figure 7 h-i) were mainly found in IC and AAF (see also Figure S6). Neurons sensitive to increases in pupil size (#1, 3, 6, 7, 8, 9, 11) (Figure 7g, i) were instead found across all auditory regions (see also Figure S6). Neurons whose response were enhanced by the presence of reward (#7-8, Figure 7f, i) were generally more abundant in subcortical regions, while those whose responses were suppressed by reward (#10, Figure 7 f, i) were mainly found in A1 (Figure 8b) (these differences are probably masked by cluster heterogeneity in each anatomically identified subpopulation in Figure S6).

In summary, we showed that highly diverse task-induced modulations influence responses of multiple cell-classes independently of their anatomical location along the auditory pathway, their laminar position in cortex, or their cortical cell type (regular or fast-spiking neurons).

## DISCUSSION

The processing of sensory stimuli varies depending on behavioral and cognitive states. Task-engagement modulation of auditory neurons has been investigated by using a variety of tasks mainly in A1 (Atiani et al., 2009; Bagur et al., 2018; Carcea et al., 2017; David et al., 2012; Francis et al., 2018a; Francis et al., 2018b; Fritz et al., 2003; Fritz et al., 2005, 2007; Kuchibhotla et al., 2017; Otazu et al., 2009) or higher cortical areas (Atiani et al., 2014; Elgueda et al., 2019). With a few exceptions (Otazu et al., 2009; Slee and David, 2015), response modulations in IC, MGB, and AAF have instead been largely overlooked. Here, we give a comprehensive description of the functional changes occurring along the auditory pathway when mice actively engage in a tone-detection task rather than passively hearing sounds. We used extracellular electrophysiological recordings to study and directly compare neuronal modulations in IC, MGB, and the two primary auditory cortices A1 and AAF. In alignment with previous reports, we found that engaging in an auditory task can induce a variety of response modulations ranging from suppression to facilitation in IC, MGB, and A1, and showed that this is also true for AAF. To our surprise, changes were more pronounced in IC, the earliest level in the hierarchy of the areas that we recorded from. Previous studies did not detect such difference, perhaps because of limited neuronal sampling (Slee and David, 2015).

Previous studies on task-induced modulations coarsely classified them as either suppressive or facilitative (Atiani et al., 2014; Atiani et al., 2009; Bagur et al., 2018; Carcea et al., 2017; David et al., 2012; Francis et al., 2018a; Francis et al., 2018b; Fritz et al., 2003; Fritz et al., 2005, 2007; Kuchibhotla et al., 2017; Otazu et al., 2009; Slee and David, 2015). The large number of recorded units and the high temporal resolution provided by electrophysiological recordings have allowed us to explore the patterns of task-engagement modulations at a much finer level. Clustering analysis of the activity changes found from passive to active listening revealed functionally distinct patterns of modulation. The changes in activity were segregated based on their time-course: they displayed enhanced or suppressed activity either only at the level of onset auditory responses (∼1/4 of the neurons) or also at the level of later response phases (∼1/2 of the neurons). We showed that enhancing onset modulations were associated with large multiplicative gain changes accompanied by stable tuning width, displaying increased responses to all tones. These modulations were absent in miss trials and therefore correlated with behavioral performance, suggesting that in the task used here such frequency unspecific facilitation may support perceptual changes improving sound detection. In contrast, suppression of onset responses in the active phase was less tied to performance and reflected divisive gain changes and narrower tuning curves. Such modulations are consistent with sparser responses in the task-engaged state that at the population level could help to discriminate between tone frequencies (Guo et al., 2017), a computation that was not required by our behavioral task. The molecular mechanisms underlying task-induced gain and tuning modulations remain to be elucidated. They could include changes in spiking thresholds or membrane conductance, changes in the balance of excitatory and inhibitory synaptic inputs, or changes in network activity such as GABAergic inhibition (Chance et al., 2002; Ferguson and Cardin, 2020; Priebe and Ferster, 2002; Zhou et al., 2014).

Previous studies on the effects that task-engagement has on the auditory system mainly adopted auditory discrimination paradigms that triggered selective attention: despite considerable variability in the modulation of single neurons, the observed changes were often suppressive (Atiani et al., 2009; David et al., 2012; Kuchibhotla et al., 2017; Slee and David, 2015). Our task is different from discrimination as it does not require animals to pay attention to one or two specific frequencies, nor does it distinguish stimuli with different auditory components. Rather, our mice had to attend to any change in their auditory environment. As we did not observe a striking prevalence of suppressive modulations, the divergent results support the hypothesis that the brain implements different neural computations to perform tone detection and discrimination tasks (Guo et al., 2017).

Interestingly, we found little difference between task-induced modulation in cortical and subcortical structures. On the one hand, this finding is consistent with previous results in IC (Slee and David, 2015) and with large-scale neural recordings that also found engagement modulation across cortical and subcortical areas during a visual task (Steinmetz et al., 2019), supporting the notion that the stations preceding cortical areas are not mere sensory relays, but can be actively involved in internal state-induced changes in sensory processing. On the other hand, our finding differs from previous studies showing that MGB is less modulated than ACx (McGinley et al., 2015; Otazu et al., 2009; Williamson et al., 2015); the structure of the task is likely to be the reason behind these differences as well. It is becoming evident that ACx is essential for difficult tasks but dispensable in easier ones (Ceballo et al., 2019; Christensen et al., 2019), and the simplicity of our behavioral paradigm is likely to have unveiled task-induced modulations that already take place in early sensory processing without the need of the more abstract stimulus representations that develop in cortex (Atiani et al., 2014). Despite the substantial similarities of task-induced modulations in cortical and subcortical auditory areas, we also found some interesting differences. In cortex but not in IC and MGB, we found a delay in tone-evoked responses in the active phase. Moreover, a difference in latency between suppressive and enhancing onset modulations was only found at the cortical level. These results may implicate the existence of neuronal processes taking place specifically in cortex such as, for example, intracortical/top-down or neuromodulatory mechanisms (Fritz et al., 2010; Goll et al., 2015; Kato et al., 2017; Winkowski et al., 2013; Winkowski et al., 2018). In addition, the latency of task-induced onset modulations revealed that they first emerge in IC and MGB, rejecting the hypothesis that subcortical modulations are only found as a consequence of cortical feedback and suggesting that subcortical auditory stations also take part in context-dependent perceptual shifts. Since we only recorded from well-trained animals that were already experts in performing the task, it is also possible that cortico-thalamic or cortico-collicular projections are needed for the development of subcortical modulations during learning (Xiong et al., 2009; Zhang and Suga, 2005; Zhang and Yan, 2008). Further experiments monitoring the activity of subcortical neurons or disrupting cortical feedback during learning will be needed to test this hypothesis.

The IC and the MGB both contain regions that are part of the lemniscal pathway (central nucleus of IC and ventral MGB) and regions that belong to the non-lemniscal pathway (dorsal and lateral nuclei of IC and medial and dorsal nuclei of the MGB). In this study we did not differentiate between these regions, and we did not find a clear bimodal distribution of either onset latency (Figure 2f-g) or tuning width (data not shown). It is possible that subcortical nuclei belonging to the non-lemniscal pathway are more modulated by task-engagement than are central nuclei (Komura et al., 2001), but we only found very weak correlations between absolute modulations amplitudes and either onset latency or tuning width (Figure S7). Future studies with more precise targeting will be needed to establish if this is indeed the case.

In addition to neurons in which task-engagement only induced modulations of the onset auditory response, half of the neurons we recorded also displayed late modulations. We showed that such late modulations were likely induced by licking, arousal, reward or combinations of them. The lick responses we observed are unlikely to reflect the well-described input from motor to auditory cortex previously shown to be suppressive (Nelson and Mooney, 2016; Nelson et al., 2013; Schneider and Mooney, 2018), but rather cholinergic inputs from the basal forebrain that has been shown to excite auditory cells during mouth movements (Nelson and Mooney, 2016). It remains to be elucidated if this projection is indeed involved, if it is the only one that causes the responses we observed, and if the same mechanism also causes subcortical licking responses. Arousal is known to affect cortical auditory neurons at the level of membrane potential, evoked response, response variability and correlation, as well as tuning (Lin et al., 2019; McGinley et al., 2015). Reward-related signals have been described in IC, non-lemniscal MGB nuclei and ACx (Brosch et al., 2011; Gruters and Groh, 2012; Guo et al., 2019; Komura et al., 2001; Metzger et al., 2006). Thus, our results confirm previous studies demonstrating arousal influences on auditory neurons and reward encoding in the auditory system. They additionally show that arousal and reward sensitivities are confined to a subset of neurons only rather than being widely distributed across units.

Besides the potential mechanisms described above, neuromodulation certainly plays a key role in many of the task-induced modulations we found, as well as during learning of the sensory-motor associations necessary for performing the task (Jacob and Nienborg, 2018; Kuchibhotla et al., 2017; Lee and Dan, 2012; Thiele and Bellgrove, 2018). In a variety of species, IC, MGB and ACx all receive extensive and diversified neuromodulation including cholinergic, dopaminergic, serotonergic and noradrenergic inputs. Further investigation will provide insights into the detailed neuromodulatory effects giving rise to the task-dependent modulations that we observed.

Taken together, our results show that the transition from passive hearing to active listening induces highly diverse patterns of functional modulation across the auditory pathway. They highlight the presence of two broad classes of neurons displaying different forms of task-induced modulations: one mainly influenced at the level of the onset auditory response, and the other being also modulated by licking, arousal and reward. This reveals the presence of functionally specialized neural subpopulations, implementing either only attentional modulation of auditory processing or also integrating information about other internal variables. The challenge for future studies is to discover the molecular or circuit mechanisms underlying the heterogeneous effects that task-engagement has on neural activity, and to understand the function that these modulations have during active listening.

## ACKNOWLEDGMENTS

This work was supported by the Swiss National Science Foundation (ERC Transfer grant to T.R.B). We thank Mari Nakamura for assistance with experiments and Magdalena Solyga, Jan Gründemann, Catherine Perrodin and Samuel Solomon for helpful comments and discussions on this article.

## AUTHOR CONTRIBUTIONS

GDF and TRB designed the study, interpreted data, and edited the manuscript. GDF performed experiments, analyzed data and wrote the manuscript. TRB acquired the funding, supervised the project and reviewed the manuscript.

## METHODS

### Animals

All experimental procedures were performed in accordance with Basel University animal care and use guidelines and were approved by the Veterinary Office of the Canton Basel-Stadt, Switzerland. Mice were derived from crossing a PV-Cre knock-in line with C57BL/6J background (JAX stock number 017320, Jackson Laboratories, ME, USA) with a ChR2-floxed Ai32 line (JAX stock n. 024109, C57BL/6J background). For all the experiments we used 12 male mice (aged 5 weeks (w) 2 days (d) - 6w6d at the start of the experiments and 8w5d - 11w4d at the end of the experiments). Water was given *ad libitum*, while food intake was manually regulated by the experimenter from the day before the start of the training phase. The animals’ weight was checked daily and maintained above 80% of the weight before food restriction. Mice were single-housed from the start of food restriction, under a 12:12 h light/dark cycle. Experiments were performed in the light phase.

### Surgical procedures

#### Headplate implant

Anesthesia was induced with Isoflurane in O_2_ (4% induction, 1.2 to 2.5% maintenance), and local analgesia was provided with subcutaneous injection of bupivacaine/lidocaine (0.01 mg/animal and 0.04mg/animal, respectively). During surgery, the depth of anesthesia was monitored by breathing rate and absence of pinch withdrawal reflex. Body temperature was maintained at 37 °C via a heating pad (FHC, ME, USA) and lubricant ophthalmic ointment was applied on both eyes. A custom-made metal head-post and a ground screw were fixed to the skull with dental cement (Super-Bond C&B; Sun Medical, Shiga, Japan). The portion of skull above the target recording site was left free from cement, and covered with a thick layer of Kwik-Cast Sealant (WPI, Sarasota, FL, USA) to protect it from external agents. Post-operative analgesia was provided with an intraperitoneal injection of buprenorphium (0.1 mg/kg). The animals recovered for at least 3 days before starting food deprivation and behavioral training. *Craniotomy*. Once animals were trained in the behavioral task, a craniotomy was performed over the region of interest. For recordings in ACx and MGB, anesthesia was induced with an intraperitoneal injection of ketamine/xylazine (80 mg/kg and 16 mg/kg, respectively) and maintained with supplementary doses of ketamine (45 mg/kg) as needed. A craniotomy (∼2.5×2.5mm) was performed above ACx or MGB, and the brain was covered by silicon oil to prevent drying. A multi-channel extracellular electrode was then used to determine the location of A1 based on functional tonotopy (caudo-rostral increase in preferred frequency), or of MGB based on the presence of auditory responses. Once A1/MGB had been located, the electrode was removed and the brain was covered by a thick layer of Kwik-Cast Sealant. For recordings in IC, animals were anesthetized with isoflurane in O_2_ (4% induction, 1.2 to 2.5% maintenance), and a craniotomy was performed above IC. The brain was then covered by a thick layer of Kwik-Cast Sealant. At the end of the surgical procedure, animals were returned to their home cage and allowed to recover until the subsequent day.

### Auditory task

Each session of the tone-detection task consisted of two passive blocks flanking one active block. Each block consisted of 150 trials. In each trial, a 1 s long pure tone of randomized frequency was presented (15 frequencies logarithmically spaced from 4-45 kHz, 65 dB SPL, 10 repetitions per frequency). In the active phase, licking within a 2 s long reward window from tone onset resulted in the delivery of a drop of vanilla soya milk as a reward (hit trial), while in case of no licking no reward was delivered (miss trial). In the passive phase, both hit and miss trials resulted in the absence of reward. Inter-trial intervals (ITI) were of random length (3-7 s). If mice licked during the ITI the timer was reset, resulting in longer periods before the next stimulus was delivered. The animals therefore had to quietly wait for a few seconds before a tone was presented. Timer resets during ITIs also served to minimize random exploratory licking behavior. Hit trials in the passive phase were not punished so that licking, or absence of it, was completely voluntary. The switch from the first passive to the active phase was cued with the release of a drop of reward that mice readily licked. The switch from the active to the second passive phase was uncued. *Behavioral training*. After the headplate implant, mice were food restricted to maintain their body weight around 80-85% of their baseline weight. Water was given ad libitum. Behavioral training started 1 day after initiation of food deprivation and was performed daily, once per day. Animals first underwent a habituation phase (1-8 days, mean 3.7 ± s.d. 0.5) during which a drop of reward was delivered if the animal licked correctly during the reward window (0-2 s after tone onset), or at the end of the reward window otherwise. As soon as the experimenter noticed an increase in performance indicating that mice had associated the auditory cue and the reward (an increase of licks in response to tones within the reward window), training was switched to the active phase (2-5 days, mean 3.3 ± s.d. 1). Once the performance in the active phase was considered by the experimenter sufficiently high and relatively stable across das, the first and second passive phases before and after the active were introduced (4-11 days, mean 5.2 ± s.d. 1.9). Once the experimenter considered the discrimination between active and passive phase sufficiently good and stable across days, a craniotomy was performed over the region of interest and recordings started.

### Pupil tracking

Mice were video monitored with a high-resolution infrared-sensitive camera (BFS-U3-16S2M-CS, FLIR Systems) focused on the eye contralateral to the recording site through a zoom lens (TCL 1216 5MP, ImagingSource, Charlotte, NC, USA). Bright uniform illumination of the eye was achieved by using two red LED lights (50668 Micro Light LED, Büchel). A third lamp in the setup provided dim white illumination to keep pupil size at intermediate levels. Frame acquisition was triggered at 30 Hz by a TTL pulse sent by the system also delivering sounds (RZ6, Tucker Davis Technologies, FL, USA). Storage of TTL timestamps allowed for post-hoc synchronization of video frames, sound stimulation, and neural recordings. The pupil was tracked using the deep learning software package DeepLabCut(Nath et al., 2019). The experimenter manually labeled the top, right, bottom and left borders of the mouse’s pupil in 259 frames extracted from videos of 8 animals, that were then used to train the network. Based on manual inspection of the extracted pupil positions’ accuracy, only extracted data with a confidence level higher than 0.01 were kept. For each frame, we estimated the pupil size as the radius of a circle fit to the available data points. Outliers (e.g. unrealistically large pupil sizes) were identified as values larger than the average pupil size + 5 times the interquartile range and removed. Only pupil traces in which less than 10% of the data points were unavailable were kept for further analysis. Missing data were filled using linear interpolation.

### Tracking of licking behavior

To track licking behavior, the licking spout was attached to a piezo detecting its movement. The voltage output from the piezo was sent to the behavioral control and data collection system (RZ6, Tucker Davis Technologies, FL, USA) where a threshold was set to detect large piezo movements corresponding to licks. The threshold was manually adjusted at the beginning of each behavioral session, and timestamps of threshold crossings were stored for offline analysis. Licking traces were calculated offline by binning the lick timestamps using the timestamps of the TTL sent to the video camera for frame acquisition (30 Hz, 0.0333 s bin width). Pupil size and licking traces were therefore synchronized. Lick traces were then smoothed using Hann window kernel filtering of 5 samples (0.1667 s).

### Neurophysiological recordings

Recordings were performed in awake, behaving mice (IC = 630 units from 4 mice: 12, 234, 231, 153 units per animal; MGB= 659 units from 3 mice: 143, 140, 376 units per animal; AAF = 1445 units from 5 mice: 64, 487, 363, 222, 309 units per animal; A1 = 1151 units from 6 mice: 136, 102, 228, 314, 292, 79 units per animal). Mice were head-fixed in a sound-attenuating chamber (modified MAC-2 chambers, Industrial Acoustics Company Nordics) in a cardboard tube in front of a licking spout. The Kwik-Cast layer was removed, and the exposed brain was covered with silicon oil. Multi-channel extracellular electrodes (Neuronexus, MI, USA. IC: 64 channels, A4×16-5mm-50-200-177-A64 in 1 session; 32 channels, A1×32-5mm-50-177-A32 in 3 sessions; A1×32-5mm-25-177-A32 in 17 sessions; MGB and ACx: 64 channels, A4×16-5mm-50-200-177-A64 in all sessions) were slowly lowered into the brain orthogonal to the surface with a motorized stereotaxic micromanipulator (DMA-1511, Narishige, Japan) (mean depth from pia ± s.d., µm. IC: 1465±215; MGB: 3519±672; AAF: 1065±56; A1: 939±25). At the end of the recording session, the electrode was removed, the brain covered with a thick layer of Kwik-Cast Sealant, and the animal returned to its home cage until the next day and recording session (mean 5.75 ± s.d. 1.42 sessions/mouse). The electrode was positioned differently day by day in each animal to maximize the number of recorded units. The analog signal was amplified, digitized and acquired at 24414Hz (RZ2 Bioamp processor, Tucker Davis Technologies, FL, USA). Putative single and multi-units were identified off-line using KiloSort(Pachitariu et al., 2016) (CortexLab, UCL, London, England) followed by manual inspection of spike shape and signal-to-noise ratio, and auto- and cross-correlograms using Phy (CortexLab, UCL, London, England). Both single and multi-units were retained for analysis.

### Auditory stimulation

Sounds were generated with a digital signal processor (RZ6, Tucker Davis Technologies, FL, USA) at 200 kHz sampling rate and played through a calibrated MF1 speaker (Tucker Davis Technologies. FL, USA) positioned 10 cm away from the mouse’s left ear. Stimuli were calibrated with a wide-band ultrasonic acoustic sensor (Model 378C01, PCB Piezotronics, NY, USA). *Tonotopy measurements:* before starting a recording during behavior, we played pure tones (50 ms duration, randomized inter-stimulus-interval distributed between 500 and 1000 ms, 2 repetitions, 6 or 0.01 ms cosine on/off ramps) varying in frequency from 4 to 48.5 kHz in 0.1 octave increments and in level from 0 to 80 dB SPL in 5 dB increments. When necessary, online analysis allowed examination of the tonotopy. *White noise:* sounds of 50 ms in duration, inter-stimulus-interval 500 ms, bandwidth of 1 to 64 kHz, 250 repetitions. We repeated the measurements at 60, 70 and 80 dB in most animals (in some just 60 and 80 dB). *Passive and active behavioral blocks:* pure tones of 1 s duration, randomized inter-stimulus-interval between 3 to 7 sec, 15 frequencies logarithmically spaced from 4-45 kHz and randomly played, 10 repetitions per frequency, 4 ms cosine on/off ramps, 65 dB SPL.

### Data analysis

Offline analysis was performed in MATLAB (release R2018a. Mathworks, MA, USA).

#### Identification of cortical layers

To identify the cortical layers, we delivered 50 ms bursts of broadband noise and performed current source density analysis of local field potentials (LFP) responses (Figure 1b). We extracted LFP by down-sampling the raw voltage traces to 1kHz and low-pass filtering (<300Hz) with an eighth order Chebyshev Type I filter. We then performed current source density analysis on the data from each electrode shank as described in Pettersen et al., 2006(Pettersen et al., 2006), using an adapted version of the CSD_plotter toolbox function ‘my_standardCSD’ (https://github.com/espenhgn/CSDplotter). Channels located in thalamo-recipient layers (input layers) were then manually identified as the ones belonging to the largest short onset current sink(Guo et al., 2017; Natan et al., 2015; Sakata and Harris, 2009; Schaefer et al., 2015; Szymanski et al., 2009; Szymanski et al., 2011). Channels located above the deepest sink were classified as located in superficial layers, those below as located in deep layers. Each unit was classified as belonging to superficial, input or deep layers based on the channel in which its spike amplitude was the largest.

#### Identification of cortical subclass

We classified cortical units as putative regular-spiking or fast-spiking (fast-spiking units are a subpopulation of parvalbumin-positive cortical interneurons). For each unit, we extracted the peak-to-trough duration (p2t) of the average spike shape. Based on the bimodal distribution of p2t across the population (Figure 1c), we defined units with a p2t < 0.5 ms as fast-spiking(Lima et al., 2009; Nowak et al., 2003).

#### Responses to tones

PSTHs were calculated for each trial from raw spike timestamps binned at 1ms bin width and smoothed using Hann window kernel filtering of 10 samples (10 ms). We then calculated an average PSTH across all trials in P1 (PSTH_P1) or P2 (PSTH_P2), and across hit (PSTH_Ahit) or miss (PSTH_Amiss) trials in A.

#### Onset latency of responses

Calculated as the earliest time point when two consecutive average PSTH bins in the first 0.05 s after tone onset exceeded with 2 STD the spontaneous rate (−0.5 to 0 s).

#### Task-engagement modulation

To quantify response modulations, average PSTHs were normalized in each unit to the maximum rate across PSTH_P1 and PSTH_Ahit (Figure S2). We then calculated a ‘modulation PSTH’ as the difference between either PSTH_Ahit, PSTH_Amiss or PSTH_P2 and PSTH_P1, so that values higher than 0 indicate increases in response and values lower than 0 indicate decreases in response from P1 to any other condition. Modulations of the spontaneous, onset or sustained rates were calculated in each unit and each condition by averaging the relevant modulation PSTH using a window of −0.5 to 0 s, 0 to 0.1 s, or 0.1 to 1 s around stimulus onset, respectively.

#### Onset latency of modulation

The average modulation of the spontaneous rate was subtracted from the modulation PSTH. Onset latency was defined as the earliest time point where two consecutive modulation PSTH bins in the first 0.1 s after tone onset exceeded 2 STD (for positive average onset modulations) or were lower than −2 STD (for negative average onset modulations) of the spontaneous rate (−0.5 to 0 s).

#### Clustering analysis

To identify clusters of neurons characterized by specific patterns of functional modulation during task-engagement, we first performed K-means clustering analysis of the modulation PSTHs between P1 trials and hit trials in A phases (PSTH_Ahit-PSTH_P1). Initial assessment of the appropriate number of clusters involved multiple runs with an increasing number of clusters, up to 20 clusters. Manual examination of the clustering results with different amounts of clusters revealed that using more than 10 clusters led to over clustering, meaning that one or more clusters displaying very similar patterns of modulations were split into different groups. We therefore set the number of clusters to 11 to allow slight over clustering, and performed 2500 runs of K-means clustering analysis (MATLAB *kmeans* function, ‘k’: 11, ‘MaxIter’: 1000, ‘Replicates’: 10, ‘distance’: correlation). We then calculated a co-clustering proportion matrix as the percentage of runs in which each pair of units were assigned to the same cluster. Only units that were put at least >90% of the K-means runs with at least one other unit were used for further clustering. To identify neurons that were consistently assigned to the same cluster, we performed hierarchical clustering analysis on the co-clustering proportion matrix (MATLAB *linkage* function) and used the resulting tree to extract 11 clusters (MATLAB *cluster* function). As mentioned above, the use of 11 clusters resulted in moderate over clustering. We therefore aggregated the 2 clusters (n = 513 and n = 54) displaying the same pattern of functional modulations (what ends up being cluster #9, Figure 4), resulting in 10 clusters. The neurons that were excluded from the hierarchical clustering step (units inconsistently assigned to clusters) were finally added in as an 11^th^ cluster.

#### Task-induced gain and frequency tuning changes

We assessed the changes in neural excitability and frequency tuning by comparing the onset responses (0-0.1 s after tone onset) to tones of different frequencies in P1 trials and hit trials of the A phase. The mean onset responses to different frequencies of each unit in each phase were fit with a Gaussian model including an additional parameter for spontaneous rate, minimizing least square errors (MATLAB *lsqcurvefit* function). Only units in which the variance explained by the model was equal to or >60% in both P1 and Ahit were analyzed further. The tuning width was estimated as the sigma of the best fit Gaussian (σ), measured in octaves. To estimate gain changes in each unit, we used the parameters of the best-fit Gaussians to simulate the response in P1 and Ahit using 0 as preferred frequency, and then normalized across P1 and Ahit. The resulting modeled P1 responses were plotted against the corresponding Ahit responses. We then extracted the best linear fit (MATLAB *polyfit* function) to the minimum and maximum values in both conditions and used the slope as an estimate of multiplicative (>1) or divisive (<1) gain changes, and the intercept as an estimate of additive (>0) or subtractive (<0) gain changes. We chose to fit the extreme values instead of the whole set of responses because many units displayed changes in tuning width which resulted in plots with a curved pattern (see Figure 5b, # 4). Fitting the whole set of rates would have therefore resulted in inaccurate estimation of gain changes.

#### Modulation by reward, arousal and licking behavior

We only analyzed neurons recorded in sessions from which we could extract both pupil and licking behavior. To quantify if neurons were sensitive to licking behavior, arousal or reward we calculated the PSTH of each unit along the active phase by binning the spike timestamps using the timebase of the acquired video frames and licking PSTHs (30 Hz, 0.0333 s bin width) and smoothed it with a Hann window kernel filtering of 5 samples (0.1667 s). We then extracted epochs (−1 to 3 s) around hit onset (first lick), or around the onset of spontaneous licking bouts initiated away from tone presentations (that is, lick bouts had to start outside the −1 to 2 s windows around tones to be considered). Using pupil size as a proxy for arousal, epochs of licking bouts away from tones were further subdivided in those accompanied by an increase in arousal or no changes in pupil size. The change in pupil size was assessed by statistical comparison of the pupil sizes measured in the 0.5 s preceding bout initiation against those measured 0.5-1.5 s after bout initiation (Wilcoxon rank-sum test). If the pupil sizes were not significantly different, the epoch was classified as one with no change in arousal level. If the statistical test resulted in significant changes, and the larger average pupil size in the two tested windows was the one after bout onset, the epoch was classified as one accompanied by an increase in arousal. We therefore extracted 3 classes of behavioral epochs: the first characterized by initiation of licking behavior and presence of reward (and a concurrent increase in pupil size), the second by licking initiation and increase in arousal but absence of reward, and the third by licking initiation in absence of both arousal changes and reward. In each session we averaged pupil size, licking traces or neural activity of each unit across epochs belonging to the same class, and then normalized across epochs. We finally quantified the neural response of each unit in different epochs by averaging its activity within a window (0-0.4 s) that maximized the similarity of common features among epochs (similar licking behavior in the three epochs, similar pupil increase between hit trials and licking bouts with arousal changes).

### Statistical analysis

Statistical tests were performed in MATLAB (release R2018a. Mathworks, MA, USA). We used non-parametric Wilcoxon rank-sum and Wilcoxon sign-rank tests, and applied Bonferroni correction to adjust for the number of hypotheses (*m*) (test and *m* are specified in figure legends). To apply Bonferroni correction, we multiplied each *p*-value by the relative *m*. In figures, statistical significance is indicated by (Bonferroni corrected) \**p<0*.*05*,\*\**p<0*.*01*, \*\*\**p<0*.*001*, \*\*\*\**p<0*.*0001*. Reported correlations are Pearson’s correlation coefficients.

### Data availability

All data and code used to generate the data are available upon reasonable request to the authors.

