## Supplemental Information for "Task-induced modulations of neuronal activity along the auditory pathway"

### SUPPLEMENTARY FIGURES AND TABLES

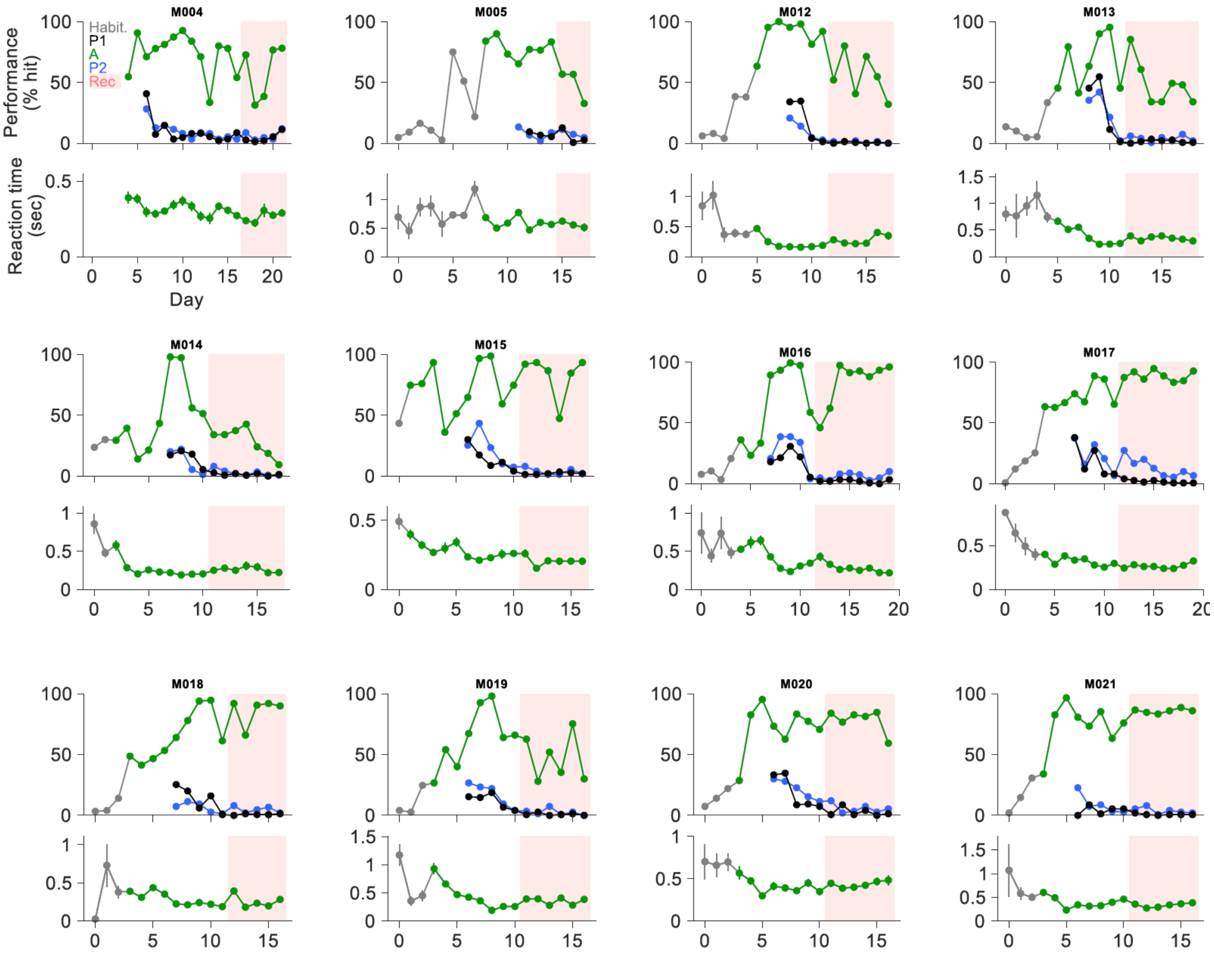


**Figure S1. Performance increases and reaction times decrease during learning. Related to Figure 1.** Learning curves for each mouse across training (white areas) and recording (blue shaded areas) days. Grey points: habituation phase. Green: active blocks (A). Black: first passive block (P1, preceding active). Blue: second passive block (P2, following active). Top: performance was measured as the percentage of correct trials over the total trials in each block. Bottom: average reaction time for hit trials during the active phase.

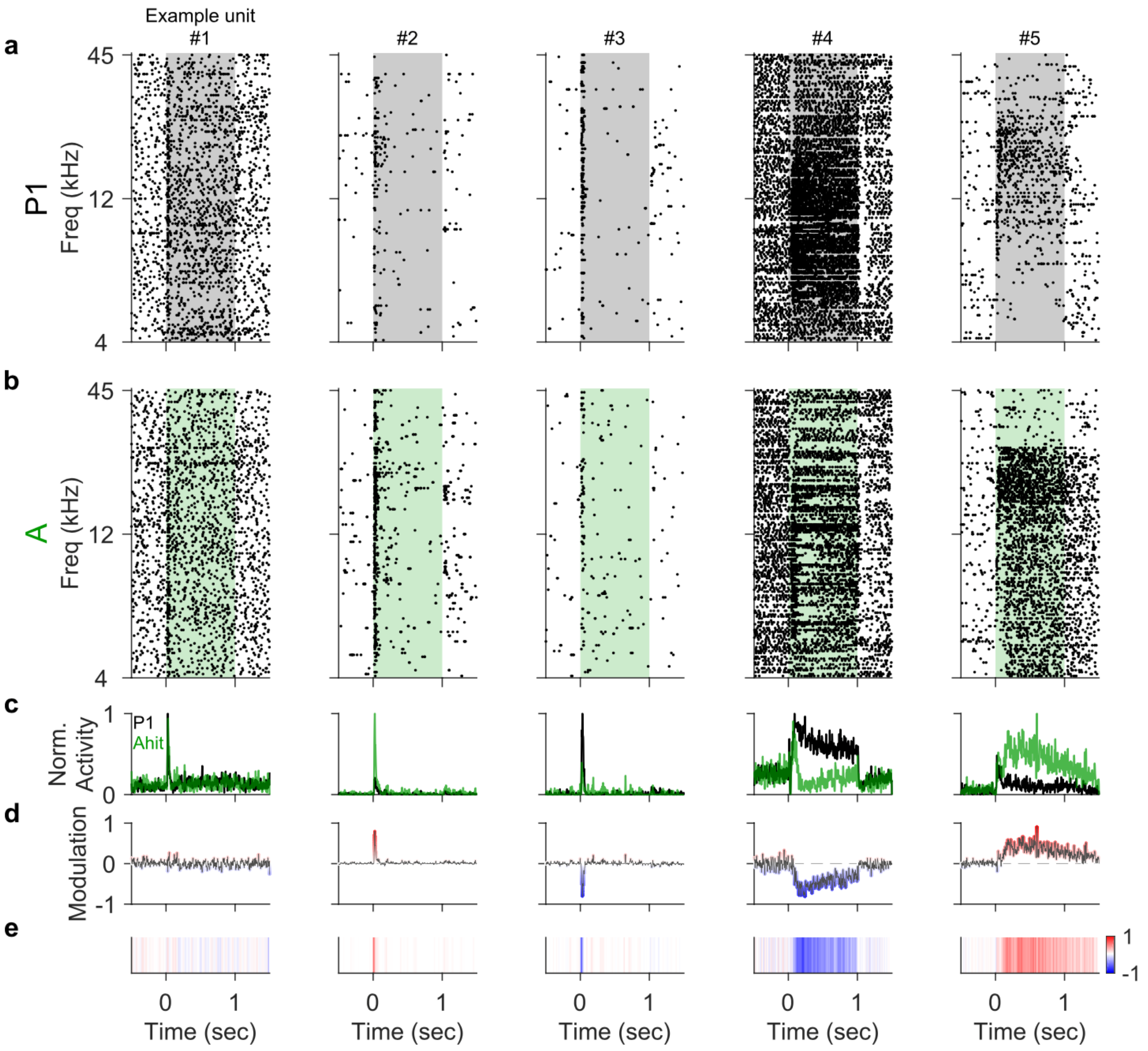


**Figure S2. Quantification of task-engagement modulation. Related to Figure 3.**
The figure illustrates the calculation of modulation PSTHs from P1 to Ahit for the 5 example units shown in Fig. 3a. Each column shows one unit (corresponding to one row in Fig. 3a). **a-b**, Raster plots showing the activity around tones of different frequency in the first passive (a) and active phase (b). Shaded areas indicate periods of tone presentation. Each dot represents one spike, each row represents one trial. Trials are ordered by tone frequency. **c**, Average PSTH across all frequencies in P1 (black) and hit trials in A (green), normalized to the maximum between the two. **d**, Modulation PSTHs depicting the change in activity around tones from P1 to Ahit in each unit (i.e. difference between black and green normalized PSTHs shown in (c)) . Red indicates an increase in activity in the active phase, blue indicates a decrease in activity. Color scale as in (e). **e**, Heatmap version of the modulation PSTH shown in (d). This visualization corresponds to one row in Fig. 3b.

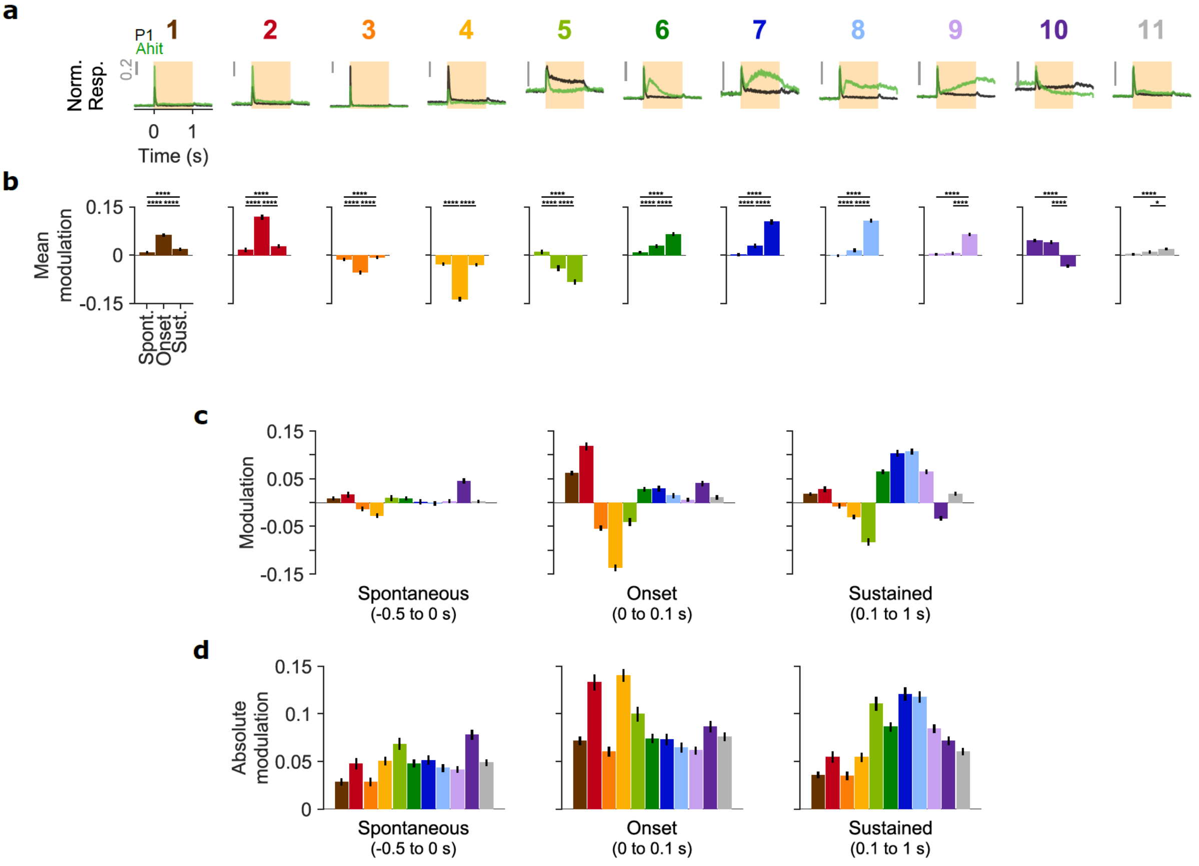


**Figure S3. Spontaneous, onset and sustained modulations in different clusters. Related to Figure 4.**
**a**, Average normalized PSTH in P1 (black) and hit trials in A (green) for each cluster. Color code as in Fig. 4. Grey vertical bar indicates 0.2 of normalized activity. **b**, Spontaneous, onset and sustained modulation across units in each cluster. Wilcoxon signed-rank test, $m$=33. **c**, **d**, Same data as (b) and Fig. 4f, respectively, highlighting comparison among clusters for spontaneous (left), onset (middle) and sustained modulations (right). Data show mean ± SEM.

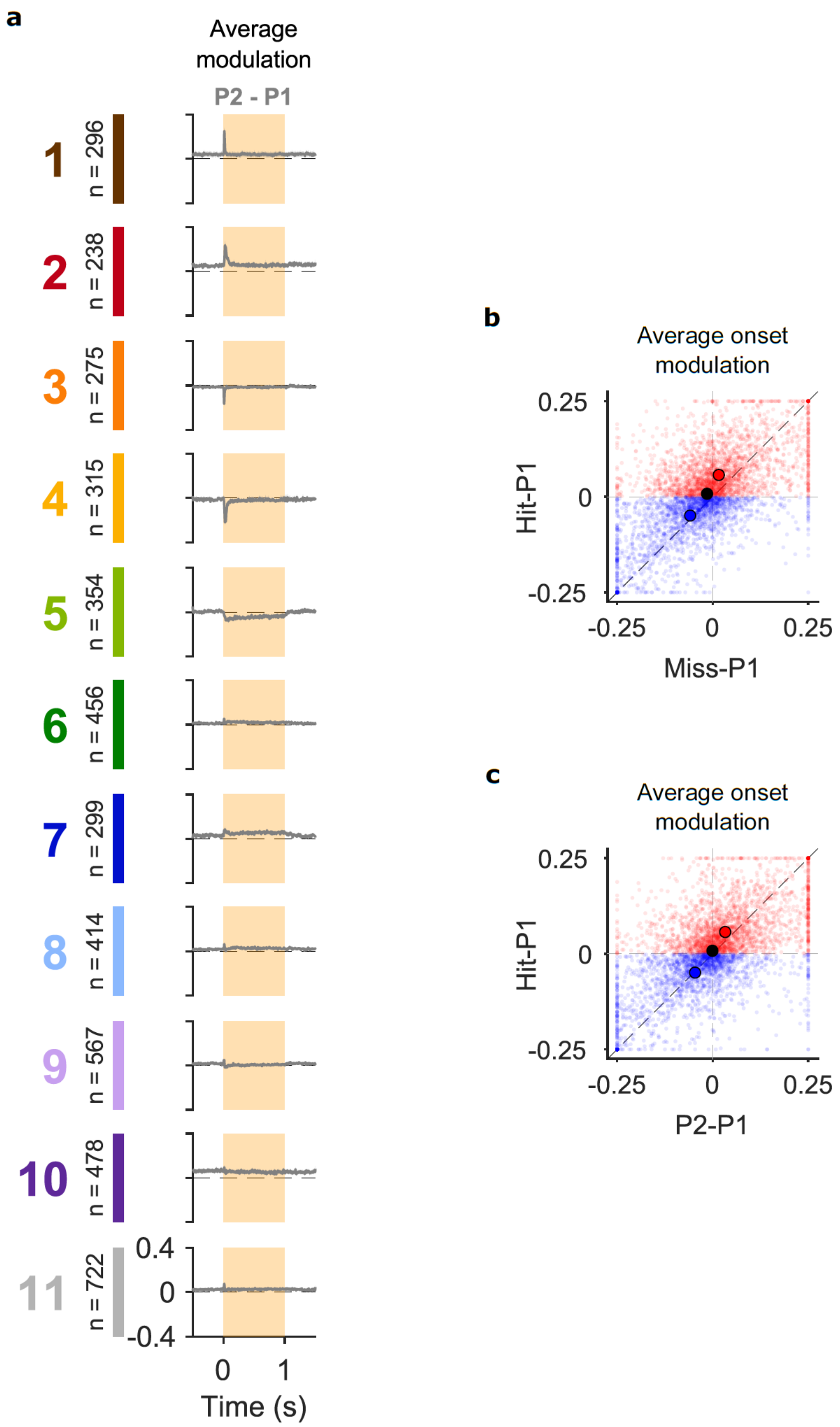


**Figure S4. Persistence of task-engagement modulation and performance specificity of onset modulations. Related to Figure 4.**

**a,** Average modulation PSTH from trials in first passive to trials in the second passive phase for neurons belonging to each cluster. Shaded areas indicate tone duration. Dashed horizontal lines indicate 0 modulation. Color code as in Fig. 4. **b**, Modulation of onset response in miss against hit trials for all units ($n = 4414$, $p < 0.0001$, Wilcoxon signed-rank test). Red: enhancing modulations in hit trials ($n = 2387$, $p < 0.0001$, Wilcoxon signed-rank test). Blue: suppressive modulations in hit trials ($n = 2000$, $p = 0.54$, Wilcoxon signed-rank test). **c**, Modulation of onset response in P2 against hit trials for all units ($n = 4414$, $p < 0.0001$, Wilcoxon signed-rank test). Red: enhancing modulations in hit trials ($n = 2387$, $p < 0.0001$, Wilcoxon signed-rank test). Blue: suppressive modulations in hit trials ($n = 2000$, $p < 0.0001$, Wilcoxon signed-rank test).

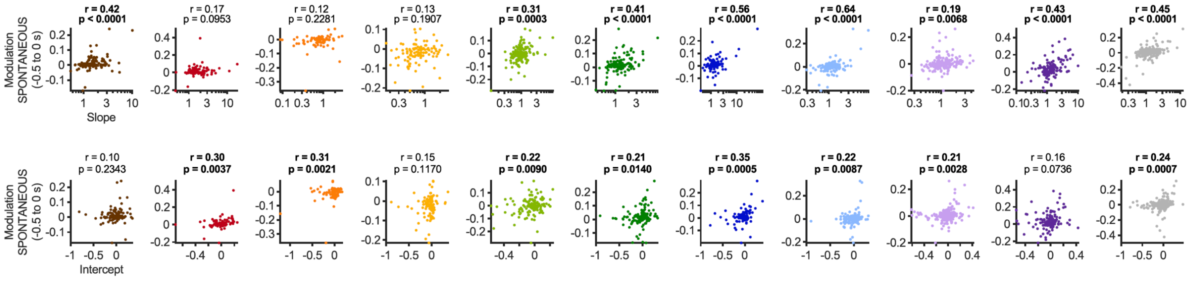


**Figure S5. Changes in gain correlate with spontaneous rate modulations. Related to Figure 5.**
Scatter plots of slope (top) and intercept (bottom) extracted from linear fits (see Fig. 5) against modulation of spontaneous rate in each cluster. Color code as in Fig. 4. Each dot is one unit. In each panel, *r* and *p* are Pearson's correlation coefficients and significance level, respectively.

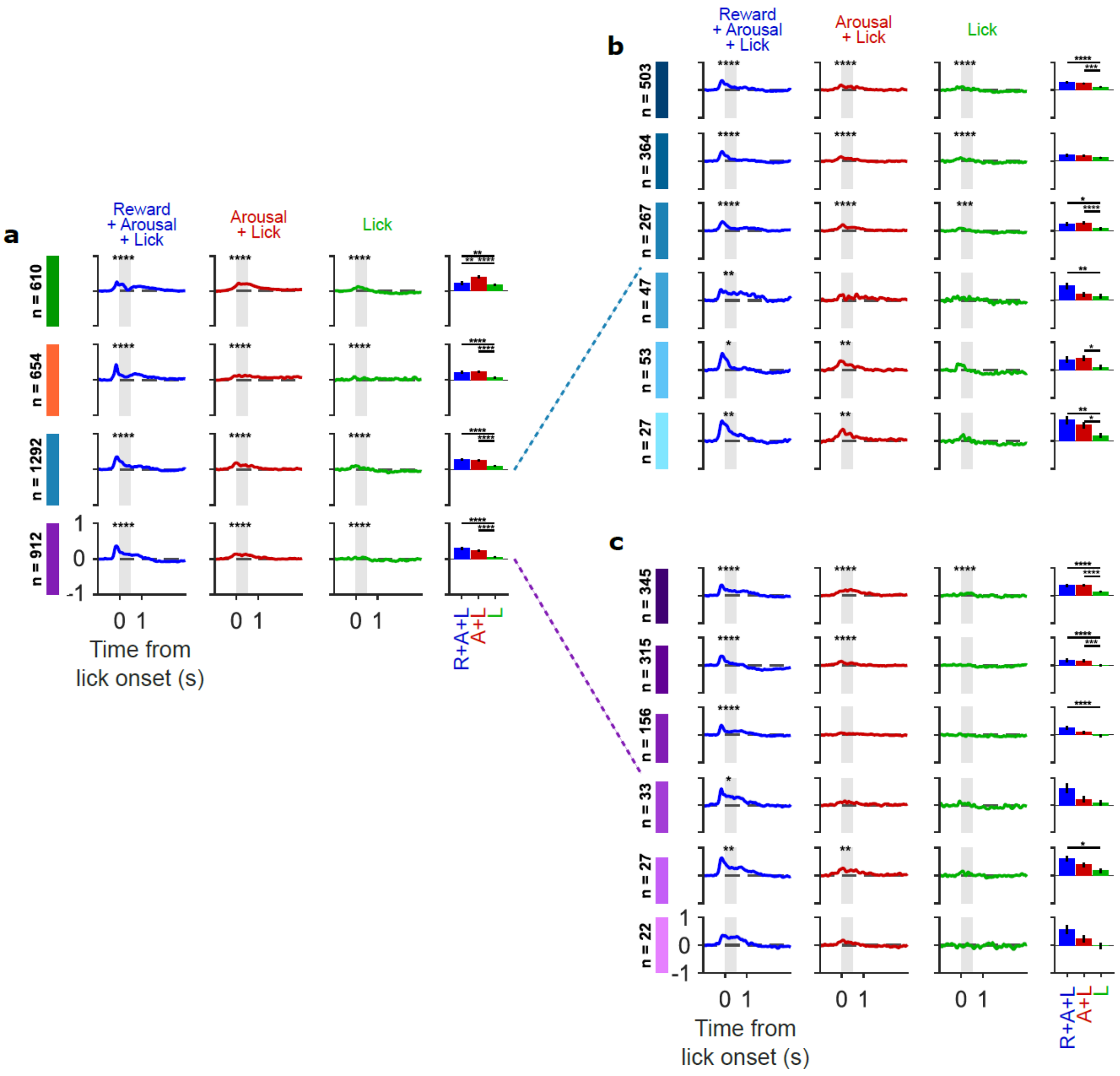


**Figure S6. Arousal, movement or reward sensitivity in different anatomical regions. Related to Figure 7.** Average normalized PSTH in epochs shown in (Fig. 7c-e) for neurons belonging to each anatomically identified area (a, one area per row), or cortical subpopulation in AAF (b) and A1 (c). Anatomical identity color code as in Fig. 2d. Blue PSTHs and bars correspond to epochs (-1 to 3 s) around the first lick in hit trials of the active phase (Fig. 7a). Red and green PSTHs and bars correspond to epochs around the onset of spontaneous licking bouts away from tone presentation accompanied by arousal increases (Fig. 7d) or not (Fig. 5e). Data show mean ± SEM. Wilcoxon signed-rank test, $m$=48.

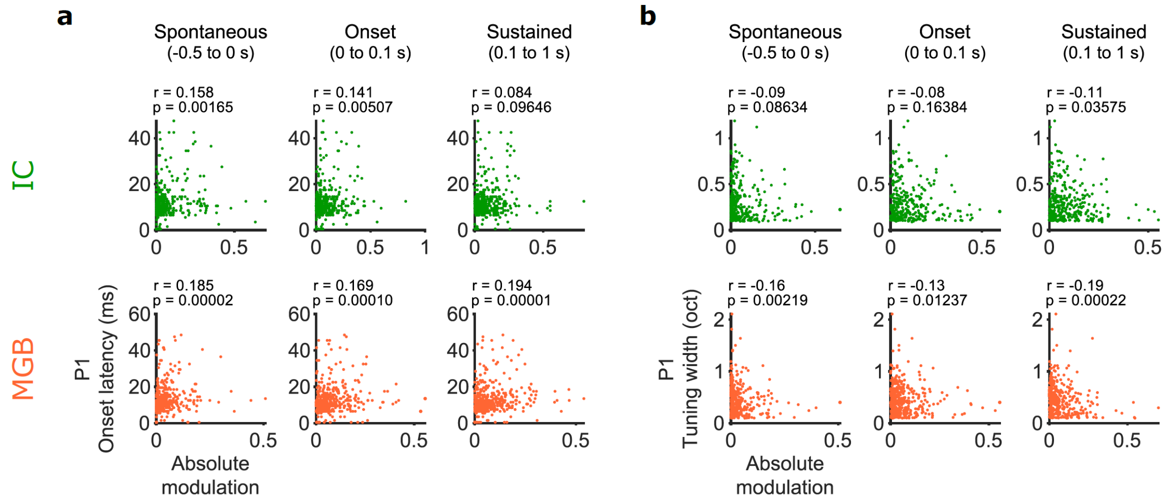


**Figure S7. Weak correlation between task-induced modulations and basic properties of tone responses in IC and MGB.** Scatter plots of absolute modulation of spontaneous (left), onset (middle) and sustained (right) modulations against onset latency (a) or tuning width (b) of responses in P1 phase in IC (green) and MGB (orange) units. Each dot is one unit. In each panel, *r* and *p* are Pearson's correlation coefficients and significance level, respectively.

**Table S1. Information about recording sessions.**

| Information about recording sessions | | | | | | | | | | | | |
|---|---|---|---|---|---|---|---|---|---|---|---|---|
| Recording Session | Animal | Area | Number of recorded units | | | | | | | Behavioral performance | | |
| | | | Total | Cell type | | Cortical layer | | | | Passive 1 | Active | Passive 2 |
| | | | | RS | FS | Sup | Inp | Deep | N/A | | | |
| **1** | gdf_m004 | A1 | 70 | 63 | 7 | 0 | 0 | 0 | 70 | 2.0 | 72.0 | 8.0 |
| **2** | gdf_m004 | A1 | 64 | 59 | 5 | 0 | 0 | 0 | 64 | 0.7 | 30.7 | 2.7 |
| **3** | gdf_m004 | unsure | 46 | 39 | 7 | 0 | 0 | 0 | 46 | 2.0 | 37.3 | 4.7 |
| **4** | gdf_m004 | AAF | 64 | 56 | 8 | 0 | 0 | 0 | 64 | 4.7 | 76.0 | 3.3 |
| **5** | gdf_m004 | unsure | 61 | 50 | 11 | 0 | 0 | 0 | 61 | 11.3 | 77.3 | 10.7 |
| **6** | gdf_m005 | A1 | 99 | 86 | 13 | 47 | 31 | 21 | 0 | 12.0 | 55.8 | 10.7 |
| **7** | gdf_m005 | unsure | 60 | 53 | 7 | 11 | 8 | 41 | 0 | 0.0 | 56.0 | 6.7 |
| **8** | gdf_m005 | unsure | 15 | 15 | 0 | 6 | 6 | 3 | 0 | 2.0 | 32.0 | 4.0 |
| **9** | gdf_m012 | A1 | 118 | 109 | 9 | 64 | 37 | 17 | 0 | 0.0 | 52.0 | 1.3 |
| **10** | gdf_m012 | A1 | 136 | 129 | 7 | 71 | 50 | 15 | 0 | 1.3 | 80.0 | 1.3 |
| **11** | gdf_m012 | AAF | 87 | 84 | 3 | 24 | 23 | 9 | 31 | 0.7 | 40.7 | 2.0 |
| **12** | gdf_m012 | AAF | 122 | 111 | 11 | 47 | 50 | 25 | 0 | 0.0 | 71.3 | 0.0 |
| **13** | gdf_m012 | AAF | 132 | 122 | 10 | 67 | 36 | 29 | 0 | 0.7 | 54.7 | 1.3 |
| **14** | gdf_m012 | AAF | 142 | 131 | 11 | 56 | 55 | 31 | 0 | 0.0 | 32.0 | 0.0 |
| **15** | gdf_m013 | A1 | 107 | 97 | 10 | 60 | 29 | 18 | 0 | 0.0 | 85.3 | 6.0 |
| **16** | gdf_m013 | A1 | 114 | 96 | 18 | 39 | 46 | 29 | 0 | 1.3 | 60.7 | 9.7 |
| **17** | gdf_m013 | A1 | 118 | 105 | 13 | 45 | 52 | 21 | 0 | 3.3 | 34.0 | 0.7 |
| **18** | gdf_m013 | AAF | 119 | 102 | 17 | 64 | 26 | 29 | 0 | 2.0 | 34.0 | 4.7 |
| **19** | gdf_m013 | AAF | 122 | 106 | 16 | 57 | 30 | 35 | 0 | 2.7 | 49.3 | 2.0 |
| **20** | gdf_m013 | AAF | 121 | 107 | 14 | 64 | 27 | 30 | 0 | 0.7 | 48.0 | 7.3 |
| **21** | gdf_m013 | IC | 12 | 0 | 12 | 0 | 0 | 0 | 12 | 0.7 | 34.0 | 2.0 |
| **22** | gdf_m014 | A1 | 117 | 110 | 7 | 35 | 27 | 33 | 22 | 2.7 | 34.0 | 8.0 |
| **23** | gdf_m014 | A1 | 97 | 88 | 9 | 39 | 42 | 16 | 0 | 0.7 | 34.0 | 4.0 |
| **24** | gdf_m014 | A1 | 100 | 93 | 7 | 36 | 44 | 20 | 0 | 2.0 | 37.3 | 2.0 |
| **25** | gdf_m014 | AAF | 97 | 91 | 6 | 30 | 45 | 22 | 0 | 0.7 | 42.7 | 0.7 |
| **26** | gdf_m014 | unsure | 130 | 116 | 14 | 63 | 34 | 33 | 0 | 2.0 | 24.0 | 3.3 |
| **27** | gdf_m014 | AAF | 125 | 114 | 11 | 55 | 44 | 26 | 0 | 0.0 | 18.7 | 0.7 |
| **28** | gdf_m014 | unsure | 85 | 78 | 7 | 15 | 46 | 24 | 0 | 1.3 | 9.3 | 0.7 |
| **29** | gdf_m015 | A1 | 84 | 72 | 12 | 15 | 34 | 16 | 19 | 1.3 | 93.3 | 4.0 |
| **30** | gdf_m015 | unsure | 68 | 62 | 6 | 21 | 31 | 5 | 11 | 2.0 | 86.7 | 1.3 |
| **31** | gdf_m015 | AAF | 83 | 69 | 14 | 23 | 39 | 21 | 0 | 3.3 | 47.3 | 1.3 |
| **32** | gdf_m015 | AAF | 139 | 122 | 17 | 65 | 47 | 27 | 0 | 2.7 | 84.7 | 5.3 |
| **33** | gdf_m015 | AAF | 86 | 73 | 13 | 21 | 34 | 31 | 0 | 2.0 | 93.3 | 2.0 |
| **34** | gdf_m016 | IC | 12 | 5 | 7 | 0 | 0 | 0 | 12 | 2.0 | 46.0 | 4.7 |
| **35** | gdf_m016 | IC | 51 | 4 | 47 | 0 | 0 | 0 | 51 | 2.0 | 62.0 | 2.7 |
| **36** | gdf_m016 | IC | 41 | 4 | 37 | 0 | 0 | 0 | 41 | 3.3 | 97.3 | 8.0 |
| **37** | gdf_m016 | IC | 30 | 4 | 26 | 0 | 0 | 0 | 30 | 3.3 | 91.3 | 8.7 |
| **38** | gdf_m016 | IC | 46 | 7 | 39 | 0 | 0 | 0 | 46 | 2.0 | 92.7 | 7.3 |
| **39** | gdf_m016 | IC | 27 | 3 | 24 | 0 | 0 | 0 | 27 | 0.7 | 88.0 | 2.7 |
| **40** | gdf_m016 | IC | 27 | 4 | 23 | 0 | 0 | 0 | 27 | 0.0 | 93.3 | 4.7 |
| **41** | gdf_m017 | IC | 18 | 14 | 4 | 0 | 0 | 0 | 18 | 4.0 | 87.3 | 27.3 |
| **42** | gdf_m017 | IC | 33 | 14 | 19 | 0 | 0 | 0 | 33 | 2.7 | 92.0 | 16.7 |
| **43** | gdf_m017 | IC | 37 | 9 | 28 | 0 | 0 | 0 | 37 | 1.3 | 86.0 | 20.0 |
| **44** | gdf_m017 | IC | 26 | 5 | 21 | 0 | 0 | 0 | 26 | 2.7 | 94.7 | 12.7 |
| **45** | gdf_m017 | IC | 34 | 8 | 26 | 0 | 0 | 0 | 34 | 1.3 | 88.7 | 6.7 |
| **46** | gdf_m017 | IC | 34 | 2 | 32 | 0 | 0 | 0 | 34 | 0.7 | 83.3 | 5.3 |
| **47** | gdf_m017 | IC | 24 | 2 | 22 | 0 | 0 | 0 | 24 | 0.7 | 84.7 | 10.0 |
| **48** | gdf_m017 | IC | 18 | 1 | 17 | 0 | 0 | 0 | 18 | 0.7 | 92.7 | 6.7 |
| **49** | gdf_m018 | IC | 43 | 2 | 41 | 0 | 0 | 0 | 43 | 0.0 | 92.0 | 8.0 |
| **50** | gdf_m018 | IC | 27 | 4 | 23 | 0 | 0 | 0 | 27 | 1.3 | 66.0 | 2.0 |
| **51** | gdf_m018 | IC | 32 | 8 | 24 | 0 | 0 | 0 | 32 | 0.7 | 90.7 | 4.7 |
| **52** | gdf_m018 | IC | 26 | 0 | 26 | 0 | 0 | 0 | 26 | 0.7 | 92.0 | 6.7 |
| **53** | gdf_m018 | IC | 24 | 1 | 23 | 0 | 0 | 0 | 24 | 1.3 | 90.0 | 2.0 |
| **54** | gdf_m019 | MGB | 24 | 2 | 22 | 0 | 0 | 0 | 24 | 0.7 | 62.7 | 3.3 |
| **55** | gdf_m019 | MGB | 37 | 3 | 34 | 0 | 0 | 0 | 37 | 2.7 | 28.0 | 1.3 |
| **56** | gdf_m019 | MGB | 16 | 0 | 16 | 0 | 0 | 0 | 16 | 0.0 | 52.0 | 7.3 |
| **57** | gdf_m019 | MGB | 18 | 8 | 10 | 0 | 0 | 0 | 18 | 0.7 | 35.3 | 0.7 |
| **58** | gdf_m019 | MGB | 21 | 5 | 16 | 0 | 0 | 0 | 21 | 1.3 | 75.3 | 2.7 |
| **59** | gdf_m019 | MGB | 34 | 11 | 23 | 0 | 0 | 0 | 34 | 0.0 | 30.0 | 0.0 |
| **60** | gdf_m020 | MGB | 79 | 4 | 75 | 0 | 0 | 0 | 79 | 8.7 | 76.7 | 2.0 |
| **61** | gdf_m020 | MGB | 23 | 2 | 21 | 0 | 0 | 0 | 23 | 4.0 | 81.3 | 7.3 |
| **62** | gdf_m020 | MGB | 17 | 3 | 14 | 0 | 0 | 0 | 17 | 0.0 | 84.7 | 2.7 |
| **63** | gdf_m020 | MGB | 20 | 2 | 18 | 0 | 0 | 0 | 20 | 1.3 | 59.3 | 5.3 |
| **64** | gdf_m021 | MGB | 54 | 35 | 19 | 0 | 0 | 0 | 54 | 2.0 | 86.7 | 5.3 |
| **65** | gdf_m021 | MGB | 78 | 39 | 39 | 0 | 0 | 0 | 78 | 0.7 | 84.7 | 8.0 |
| **66** | gdf_m021 | MGB | 37 | 16 | 21 | 0 | 0 | 0 | 37 | 0.0 | 83.3 | 0.7 |
| **67** | gdf_m021 | MGB | 66 | 21 | 45 | 0 | 0 | 0 | 66 | 0.7 | 86.0 | 4.0 |
| **68** | gdf_m021 | MGB | 78 | 27 | 51 | 0 | 0 | 0 | 78 | 0.7 | 88.7 | 2.7 |
| **69** | gdf_m021 | MGB | 62 | 35 | 27 | 0 | 0 | 0 | 62 | 0.7 | 86.0 | 2.0 |

**Table S2. Tone evoked onset latency in P1 vs Ahit in each anatomically identified population.** Statistical test was Wilcoxon signed-rank test with *m*=16.

| Tone-evoked Onset Latency | | | | | | | | |
|---|---|---|---|---|---|---|---|---|
| Anatomical Region | p-value | h | Passive 1 | | | Active Hit | | |
| | | | n | Median (ms) | SEM | n | Median (ms) | SEM |
| **IC** | 12.4 | 0 | 319 | 10.5 | 0.30 | 319 | 10.5 | 0.31 |
| **MGB** | 4.93 | 0 | 464 | 10.5 | 0.29 | 464 | 10.5 | 0.28 |
| **AAF** | 0.000000000 | 1 | 919 | 12.5 | 0.23 | 919 | 14.5 | 0.26 |
| AAF-SUP-RS | 0.0000013 | 1 | 327 | 16.5 | 0.37 | 327 | 17.5 | 0.40 |
| AAF-INP-RS | 0.000023 | 1 | 290 | 12.5 | 0.41 | 290 | 13.5 | 0.51 |
| AAF-DEEP-RS | 0.000054 | 1 | 183 | 10.5 | 0.44 | 183 | 11.5 | 0.57 |
| AAF-SUP-FS+ | 2.37 | 0 | 36 | 15 | 1.29 | 36 | 15.5 | 0.76 |
| AAF-INP-FS+ | 0.061 | 0 | 56 | 10 | 0.35 | 56 | 11 | 0.83 |
| AAF-DEEP-FS+ | 0.013 | 1 | 27 | 10.5 | 1.51 | 27 | 11.5 | 0.99 |
| **A1** | 0.0000053 | 1 | 601 | 20.5 | 0.36 | 601 | 21.5 | 0.38 |
| A1-SUP-RS | 0.000068 | 1 | 243 | 21.5 | 0.50 | 243 | 22.5 | 0.50 |
| A1-INP-RS | 0.056 | 0 | 194 | 19.5 | 0.66 | 194 | 20.5 | 0.73 |
| A1-DEEP-RS | 6.63 | 0 | 94 | 17.5 | 1.15 | 94 | 20.5 | 1.14 |
| A1-SUP-FS+ | 15.1 | 0 | 28 | 20.5 | 1.43 | 28 | 21.5 | 1.62 |
| A1-INP-FS+ | 4.03 | 0 | 29 | 13.5 | 1.59 | 29 | 15.5 | 1.84 |
| A1-DEEP-FS+ | 14.6 | 0 | 13 | 22.5 | 2.13 | 13 | 20.5 | 2.56 |

**Table S3. Tone evoked onset latency in P1 vs Ahit in each functional cluster.** Statistical test was Wilcoxon signed-rank test with *m*=11.

| Tone-evoked Onset Latency | | | | | | | | |
|---|---|---|---|---|---|---|---|---|
| Cluster | p-value | h | P1 | | | AHit | | |
| | | | n | Median (ms) | SEM | n | Median (ms) | SEM |
| 1 | 4.10 | 0 | 235 | 12.50 | 0.50 | 235 | 13.5 | 0.35 |
| 2 | 0.78 | 0 | 169 | 19.50 | 0.71 | 169 | 21.5 | 0.64 |
| 3 | 0.000000046 | 1 | 229 | 10.50 | 0.23 | 229 | 10.5 | 0.46 |
| 4 | 0.00096 | 1 | 201 | 16.50 | 0.50 | 201 | 18.5 | 0.71 |
| 5 | 0.51 | 0 | 175 | 18.50 | 0.73 | 175 | 19.5 | 0.77 |
| 6 | 0.0000023 | 1 | 302 | 11.50 | 0.42 | 302 | 12.5 | 0.48 |
| 7 | 0.031 | 1 | 143 | 14.50 | 0.64 | 143 | 16.5 | 0.79 |
| 8 | 0.010 | 1 | 289 | 12.50 | 0.44 | 289 | 13.5 | 0.45 |
| 9 | 0.099 | 0 | 344 | 13.50 | 0.41 | 344 | 14.5 | 0.44 |
| 10 | 0.000026 | 1 | 184 | 18.50 | 0.63 | 184 | 20.5 | 0.79 |
| 11 | 0.00048 | 1 | 412 | 14.50 | 0.43 | 412 | 15.5 | 0.46 |

**Table S4. Task-induced modulation onset latency in enhanced vs suppressed modulations in each anatomically identified population.** Statistical test was Wilcoxon signed-rank test with *m*=16.

| Task-induced Modulation Onset Latency | | | | | | | | |
|---|---|---|---|---|---|---|---|---|
| Area | *p*-value | h | Enhanced | | | Suppressed | | |
| | | | n | Median (ms) | SEM | n | Median (ms) | SEM |
| **IC** | 8.74 | 0 | 302 | 12.5 | 1.27 | 200 | 11.5 | 1.27 |
| **MGB** | 0.246 | 0 | 333 | 13.5 | 1.04 | 218 | 11.5 | 1.04 |
| **AAF** | 0.000004 | 1 | 532 | 19.5 | 0.87 | 635 | 14.5 | 0.87 |
| AAF-SUP-RS | 9.62 | 0 | 203 | 20.5 | 1.45 | 247 | 20.5 | 1.23 |
| AAF-INP-RS | 0.42 | 0 | 153 | 16.5 | 1.43 | 186 | 13.5 | 1.09 |
| AAF-DEEP-RS | 0.00000001 | 1 | 112 | 22.5 | 2.00 | 137 | 10.5 | 1.47 |
| AAF-SUP-FS | 12.10 | 0 | 26 | 18 | 2.96 | 19 | 16.5 | 3.37 |
| AAF-INP-FS | 0.092 | 0 | 21 | 13.5 | 4.12 | 35 | 9.5 | 1.42 |
| AAF-DEEP-FS | 0.064 | 0 | 17 | 23.5 | 6.26 | 11 | 7.5 | 7.12 |
| **A1** | 0.0088 | 1 | 451 | 20.5 | 0.81 | 421 | 25.5 | 0.81 |
| A1-SUP-RS | 0.051 | 0 | 189 | 22.5 | 1.17 | 151 | 26.5 | 1.43 |
| A1-INP-RS | 7.17 | 0 | 141 | 19.5 | 1.49 | 153 | 23.5 | 1.45 |
| A1-DEEP-RS | 3.99 | 0 | 68 | 21 | 2.49 | 87 | 24.5 | 2.46 |
| A1-SUP-FS | 0.024 | 1 | 22 | 20.5 | 3.24 | 13 | 38.5 | 6.93 |
| A1-INP-FS | 7.42 | 0 | 20 | 16 | 3.21 | 9 | 18.5 | 7.14 |
| A1-DEEP-FS | 4.18 | 0 | 11 | 13.5 | 3.92 | 8 | 17 | 5.71 |
